# Identification of novel regulators of human naïve pluripotency by means of a genetic switch utilizing the chimeric receptor G-CSFR:GP130

**DOI:** 10.1101/2025.04.09.647577

**Authors:** Tahereh Kiani, Claire Santamaria, Nathalie Doerflinger, Aikaterini Kafousi, Cloé Rognard, Ambre Bender, Michael Dumas, Michael Weber, Alexandre Bruneau, Laurent David, Pierre Savatier, Pierre-Yves Bourillot

## Abstract

This study investigated the molecular mechanisms underpinning the transition from primed to naive pluripotency in human pluripotent stem cells (hPSCs) by utilizing a dual genetic switch combining hormone-dependent STAT3-ER^T2^ and a chimeric GCSFR:gp130 receptor. Upon activation with tamoxifen and G-CSF, hPSCs demonstrated upregulation of naive markers, and sustained self-renewal independent of LIF. Transcriptomic analyses revealed distinct clusters of early response genes, with a subset of 26 genes significantly enriched in the human epiblast. Knockdown experiments further delineated essential regulators that modulate differentiation and proliferation during naive reprogramming including interferon gamma inducible protein (IFI)16 and interferon-induced transmembrane (IFITM) proteins. Moreover, functional assays using mutant chimeric GCSFR:gp130 receptor highlighted the critical role of JAK kinase recruitment in orchestrating epigenomic remodeling and early gene activation. This approach establishes a robust platform to dissect the signaling dynamics governing the accession of hPSCs to naive pluripotency.

## Introduction

Over the past decade, various protocols have successfully reverted conventional human pluripotent stem cells (PSCs) to a naïve-like state, including NHSM, 3iL, Reset, 5iLA/5iLAF, 2iL, and TL2i (Chan *et al*, 2013; Chen *et al*, 2015; Duggal *et al*, 2015; Gafni *et al*, 2013; Gao *et al*, 2019; Qin *et al*, 2016: Guo, 2017 #5875; Takashima *et al*, 2014; Theunissen *et al*, 2014; Ware *et al*, 2014; Yang *et al*, 2017). All these methods incorporate human LIF in the culture medium, inducing transcriptomic and epigenomic changes that mirror those of rodent naïve cells. Across studies, an upregulation of the LIF/JAK/STAT3 pathway has been consistently reported, including increased expression of LIF receptors (LIFR and GP130/IL6ST) (Chan *et al*., 2013; Chen *et al*., 2015; Guo *et al*, 2016; Qin *et al*., 2016; Ware *et al*., 2014), enhanced STAT3 phosphorylation (Chan *et al*., 2013; Chen *et al*., 2015; Hanna *et al*, 2010), and activation of STAT3 target genes (Chan *et al*., 2013; Chen *et al*., 2015; Duggal *et al*., 2015; Guo *et al*, 2017; Guo *et al*., 2016; Hanna *et al*., 2010; Qin *et al*., 2016; Takashima *et al*., 2014; Theunissen *et al*., 2014; Yang *et al*., 2017). Functional studies further confirm the central role of the LIF/JAK/STAT3 pathway in supporting self-renewal in naïve-like PSCs. Inhibiting JAK/STAT3 signaling–either through JAK inhibitors or by reducing STAT3 activity)– promotes differentiation or growth retardation, whereas constitutive STAT3 activation sustains the undifferentiated state (Chan *et al*., 2013; Chen *et al*., 2015; Gafni *et al*., 2013; Hanna *et al*., 2010; Ware *et al*., 2014). Single-cell RNA-sequencing of human preimplantation embryos further reveals that epiblast cells express core components of the GP130 pathway, including IL6R, GP130, JAK1, and STAT3, and the negative regulator SOCS3 (Bourillot *et al*, 2019). Interestingly, while LIF itself is scarcely expressed in any embryonic lineage, its receptor is widely present, suggesting that although LIF may not be an endogenous embryonic signal, epiblast cells remain poised to respond to it. Similar patterns are observed in cynomolgus monkey embryos and their naïve-like PSCs, where preimplantation epiblasts express both naïve markers and GP130/JAK/STAT3 components, with expression declining post-implantation (Bourillot *et al*., 2019). Collectively, these findings suggest that the GP130/JAK/STAT3 signaling machinery is an intrinsic feature of the naïve epiblast and that reprogramming primed hPSCs to a naïve-like state reinstates this pathway and LIF responsiveness.

Despite these insights, little is known about the early events that occur as human PSCs reestablish GP130/JAK/STAT3 signaling during the transition to naïve pluripotency. Key questions remain: Is STAT3 activation alone sufficient to initiate the naïve program, or are additional factors required? What role does the GP130/JAK/STAT3 pathway play in the epigenomic remodeling that accompanies this transition? To address these questions, we developed a genetic switch to induce the transition from the primed to the naïve-like state in human PSCs. This system consists of two components: a hormone-dependent STAT3-ER^T2^, which is activated by tamoxifen (Chen *et al*., 2015), and a chimeric GCSFR:gp130 receptor, which dimerizes in response to G-CSF (Niwa *et al*, 1998). Using this approach, we investigated the role of GP130-associated factors in naïve-state conversion, and identified novel genes activated during the early stages of this transition. Our findings provide new insights into the mechanisms by which human PSCs acquire naïve pluripotency.

## Results

### A LIF-independent dual genetic switch for naive state conversion

To induce the transition of hESCs from a primed to a naïve-like state, we developed a genetic switch combining two components: a hormone-dependent STAT3-ER^T2^ and a chimeric GCSFR:gp130 receptor. This receptor consists of the extracellular domain of the human granulocyte colony-stimulating factor receptor (GCSFR/GR) fused to the cytoplasmic domain of the mouse gp130 signal transducer (**Fig. 1A**). In mouse ESCs, GCSFR:gp130 dimerization by G-CSF leads to STAT3 and SHP2 phosphorylation, inhibiting differentiation even in the absence of LIF (Burdon *et al*, 1999; Niwa *et al*., 1998). Similarly, in hESCs expressing this chimeric receptor (hereafter OS3-GRgp130wt), G-CSF treatment induced phosphorylation of STAT3 (Tyr 705) and of SHP2 (Tyr 542) (**Figs. 1B, S1A**). However, unlike in mESCs, G-CSF alone failed to sustain OS3-GRgp130wt self-renewal (**Fig. 1C**), consistent with previous reports that primed hESCs do not respond to LIF alone (Daheron *et al*, 2004; Sumi *et al*, 2004). Notably, adding tamoxifen (0.5 µg/ml) to activate STAT3-ER^T2^ significantly increased STAT3 phosphorylation (**Fig. 1B**). After seven days of G-CSF and tamoxifen treatment in FGF2-free conditions, OS3-GRgp130wt cells formed undifferentiated colonies (**Fig. 1C**) and expressed core pluripotency markers (*POU5F1/OCT4*, *SOX2*, *NANOG*) alongside primed markers (*CER1*, *LEFTY1*, *LEFTY2*, *ACVR2A*, and *ACVR2B*) (**Figs. 1D, S1B**). Notably, they also up-regulated naïve-associated genes, including *ESRRB*, *DPPA5*, *FGF4*, *KLF2*, *KLF4*, *KLF5*, *KLF17*, *NROB1*, *PRDM14*, *REX1*, *TBX3*, *TFE3*, *SALL4*, and *TFCP2L1*. In contrast, parental OS3 cells lacking the chimeric receptor failed to maintain pluripotency under the same conditions (**Figs**. **1C, 1D, S1B**). Importantly, neither G-CSF, tamoxifen, nor LIF alone was sufficient to sustain pluripotency in FGF2-deprived OS3-GRgp130wt cells (**Figs. 1C, 1D, S1B**). Instead, our results indicate that activation of both the GCSFR:gp130 receptor and STAT3-ER^T2^ acts synergistically to promote self-renewal and naïve marker expression. Notably, OS3-GRgp130wt cells responded similarly to LIF + tamoxifen, supporting our previous findings that this combination facilitates conversion from the primed to naive state in STAT3-ER^T2^-expressing OS3 cells (Chen *et al*., 2015). Furthermore, this process was strictly JAK-dependent.

**Figure 1:**
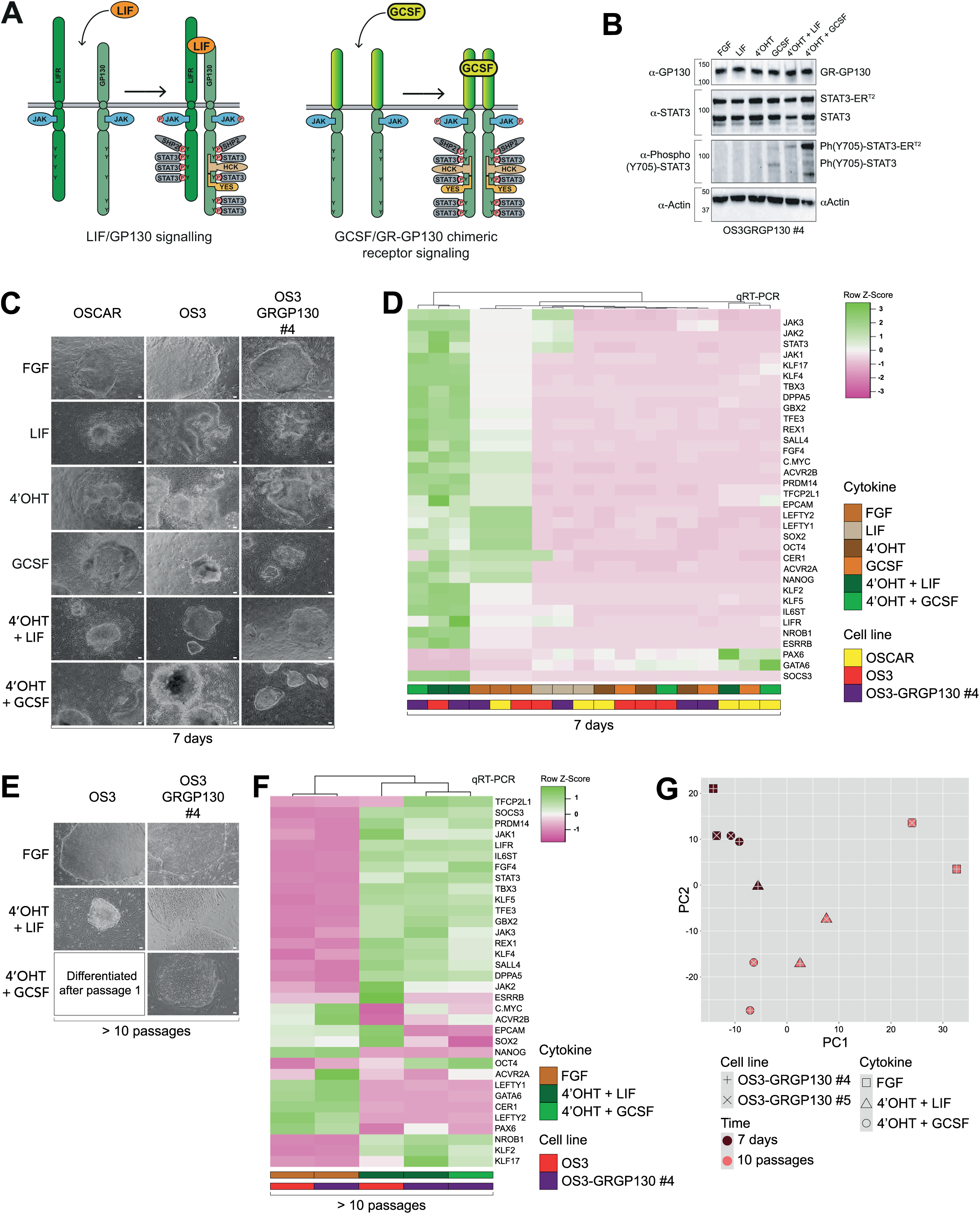
Synergistic activation of the GCSF:GP130 receptor and STAT3-ERT2 sustains self-renewal and pluripotency in FGF2-deprived OS3-GRgp130wt cells. **(A)** Schematic representation of the LIF/GP130 receptor and chimeric GCSF:GP130 receptor, combining the extracellular domain of GCSF-R and the cytoplasmic domain of GP130. **(B)** Western blot analysis of GP130, STAT3, phospho-(Tyr705)-STAT3 expression in OS3-GRgp130wt #4 cells cultured with FGF (5 ng/mL), LIF (10,000 U/mL), 4’OHT (250 nM), G-CSF (3 µM), 4’OHT (250 nM) + LIF (10,000 U/mL) and 4’OHT (250 nM) + GCSF G-CSF (3 µM). (**C**) Phase-contrast microphotography of OSCAR, OS3 and OS3-GRgp130wt #4 cells cultured with FGF, LIF, 4’OHT, GCSF, 4’OHT+LIF and 4’OHT+GCSF, for 7 days. Scale Bar, 50µm. (**D**) Hierarchical clustering and heatmap of qRT-PCR data shown in **Fig. S1B** (gene expression analysis of pluripotency (POU5F1/OCT4, SOX2, NANOG), primed pluripotency (CER1, LEFTY1/2, ACVR2A/B), naive pluripotency (ESRRB, DPPA5, FGF4, KLF2/4/5/17, NROB1, PRDM14, REX1, TBX3, TFE3, SALL4, TFCP2L1), differentiation (GATA6, PAX6) and GP130 signaling (C-MYC, IL6ST, JAK1/2/3, LIFR, SOCS3, STAT3) markers in OSCAR, OS3 and OS3-GRgp130wt #4 cells treated with FGF, LIF, 4’OHT, GCSF, 4’OHT+LIF and 4’OHT+GCSF, for 7 days) using Euclidean coefficient as a measure of distance between rows and columns and the complete linkage as a clustering method. (**E**) Phase-contrast microphotography of OS3 and OS3-GRgp130wt #4 cells cultured with FGF, 4’OHT+LIF and 4’OHT+GCSF, over 10 passages. (**F**) Hierarchical clustering and heatmap of qRT-PCR data shown in **Fig. S1C** (gene expression analysis of markers used in (D), in OS3 and OS3-GRgp130wt #4 cells cultured with FGF, 4’OHT+LIF and 4’OHT+GCSF, over 10 passages) using the same clustering methods as in (D). (**G**) Principal component analysis (PCA) of whole transcriptome data from two independent clones (OS3-GRgp130wt #4 and OS3-GRgp130wt #5) maintained in FGF, 4’OHT+LIF and 4’OHT+GCSF, for 7 days or over 10 passages.

To evaluate long-term self-renewal, OS3-GRgp130wt cells were cultured for 10 passages under CSF + tamoxifen. While control OS3 cells differentiated after the first passage, OS3-GRgp130wt cells maintained undifferentiated colonies (**Fig. 1E**). Transcriptomic analyses confirmed repression of primed markers and activation of naïve markers under G-CSF + tamoxifen, comparable to LIF + tamoxifen (**Figs. 1F, S1C**). These results suggest that GCSFR:gp130 activation by G-CSF functionally mimics LIFR activation by LIF.

To further validate our findings, we performed transcriptome analysis on two independent OS3-GRgp130wt clones (OS3-GRgp130wt#4 and OS3-GRgp130wt#5). Both clones exhibited similar self-renewal behavior under G-CSF + tamoxifen or LIF + tamoxifen (**Fig. S1D**). Principal component analysis (PCA) confirmed that these clones clustered together across different conditions, underscoring the robustness of our model (**Figs. 1G, S1E**).

### Identification of early response genes in the primed-to-naive transition

To identify early response genes activated during the transition from a primed to a naïve state, we treated two independent OS3-GRgp130wt cell clones (#4 and #5) with G-CSF, LIF, tamoxifen, and their combinations (G-CSF + tamoxifen, LIF + tamoxifen, and G-CSF + LIF + tamoxifen) for 12 and 24 hours, followed by RNA sequencing (**Fig. 2A**). No significant transcriptomic differences were observed between the two clones, allowing for pooled analysis (**Fig. S2A**). A UMAP representation revealed six distinct clusters (**Figs. 2B, 2C**): Clusters 1, 2, 4 and 5 corresponded to cells treated with single molecules for 12 and 24 hours; Clusters 0 and 3 included cells treated with G-CSF + tamoxifen, LIF + tamoxifen, and G-CSF + LIF + tamoxifen, with cluster 0 predominating at 12 hours and cluster 3 at and 24 hours (**Fig. 2D**). Differential gene expression analysis (DEGs) between LIF + tamoxifen, G-CSF + tamoxifen, and LIF + G-CSF + tamoxifen to FGF2-treated cells showed that most DEGs were upregulated in response to combinatorial treatments (**Fig. 2E, S3**). Among them, the 100 most differentially expressed genes were significantly enriched in clusters 0 and 3 compared to clusters 1, 2, 4 and 5 (**Fig. S2B**). A heatmap confirmed that these genes were specifically upregulated in combinatorial conditions, while single-molecule treatments had minimal effect (**Fig. 2F**). Notably, G-CSF + tamoxifen elicited a stronger transcriptional response than LIF + tamoxifen, suggesting superior efficiency in converting OS3-GRgp130wt cells.

**Figure 2:**
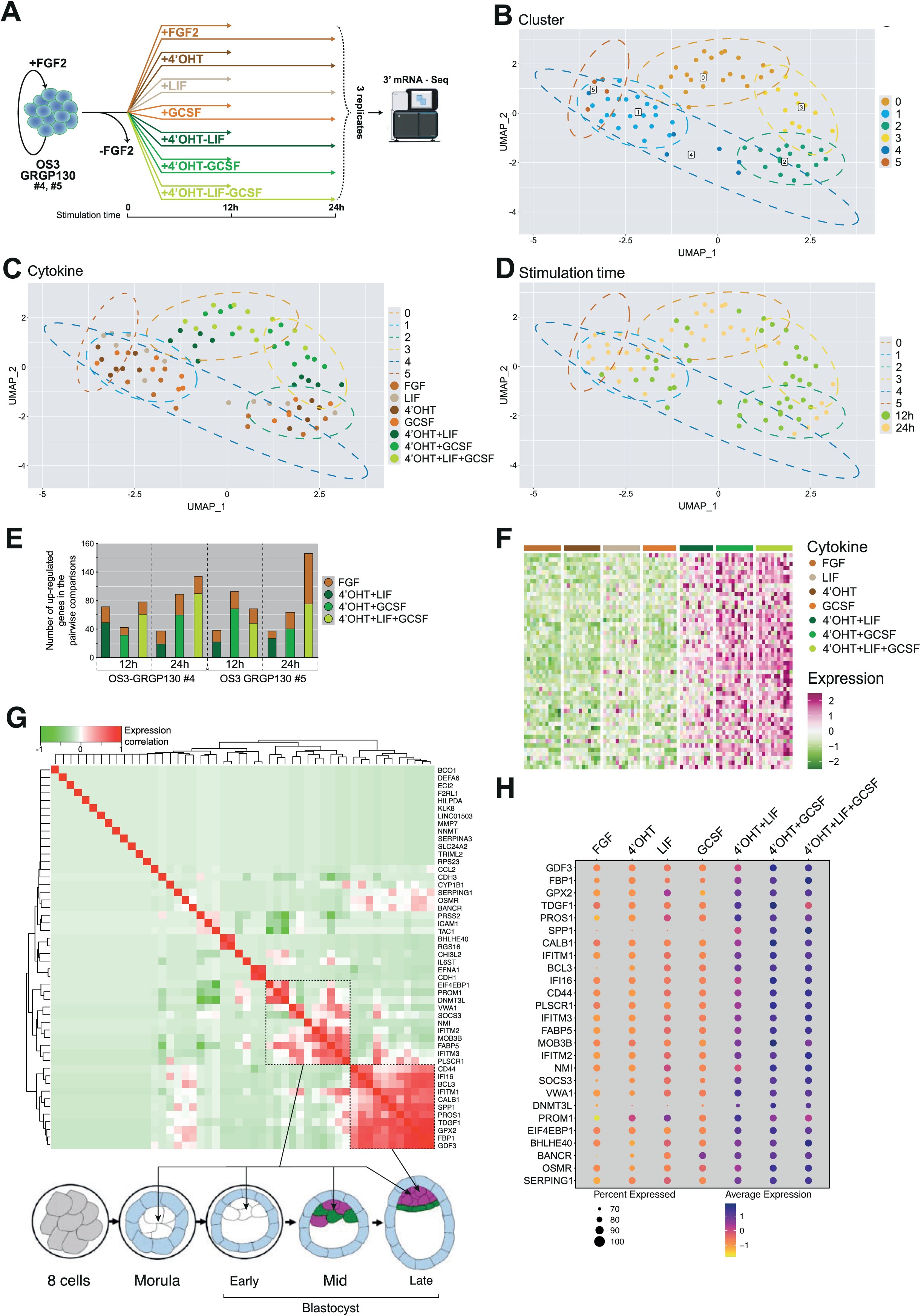
Identification of early response genes induced by G-CSF and tamoxifen in OS3-GRgp130wt cells. **(A)** Schematic representation of the experimental design: two independent OS3-GRgp130wt cell clones (#4 and #5) were treated for 12 and 24 hours with FGF, G-CSF, LIF, tamoxifen, or the combinations (G-CSF + 4’OHT, LIF + 4’OHT, and G-CSF + LIF + 4’OHT), followed by RNA sequencing. **(B)** UMAP plot from RNA-seq analysis of the samples described in (A). Samples are coloured according to the Seurat cluster. **(C)** UMAP plot coloured according to the culture conditions. **(D)** UMAP plot coloured according to the stimulation time. (E) Histogram showing the number of up-regulated genes identified in pairwise comparisons between LIF + 4’OHT, G-CSF + 4’OHT, and LIF + G-CSF + 4’OHT versus FGF2, in OS3-GRgp130wt cells after 12 and 24 hours of treatment. (**F**) Heatmap showing the expression of the 100 most differentially expressed genes identified in **Figs. 2E, S3**. Samples are grouped according to the culture conditions. Expression levels are row-wise z-transformed DESeq2-normalized counts. (**G**) Correlation heatmap showing the expression correlation between 50 DEGs in the human pre-implantation embryo. Correlation expression scores are taken from https://humanembryo-ui.bird.glicid.fr/human/PTUI.html (Meistermann *et al*., 2021). (**H**) Dot plot showing the expression of the 25 short-listed DEG in OS3-GRgp130wt cells, across the different culture conditions. Dot size represents the proportion of samples in the indicated group expressing the given gene and colour encodes the scaled average expression.

To assess their relevance in early human development, we examined the expression of these 100 DEGs in human preimplantation embryos using the dataset from Meistermann et al. (Meistermann *et al*, 2021). This analysis identified 11 genes highly enriched in the epiblast compared to the trophectoderm and primitive endoderm in blastocysts (**Fig. 2G**). Another set of 11 genes were strongly enriched in pluripotent cells from the morula to blastocyst stages. Among these 22 genes, only *TDGF1* (*CRIPTO*) (Fiorenzano *et al*, 2016) and *DNMT3L* (Wu *et al*, 2021) have been previously linked to naïve pluripotency in mice. The remaining genes encode a diverse range of proteins, including interferon-inducible proteins (*IFITM1*, *IFITM2*, *IFITM3*, *IFI16*, *PLSCR1*, *NMI*), metabolic regulators (*FBP1*, *GPX2*), and developmental factors (*PROS1*, *SENP6*, *CALB1*, *BCL3*, *CD44*, *VWA1*, *MOB3B*, *FABP5*, *PROM1*, *EIF4EBP1*). In addition to these 22 genes, we included the long non-coding RNA, *BANCR,* Oncostatin M receptor (OSMR), Serpin Family G Member 1 (SERPING1), and Basic Helix-Loop-Helix Family Member E40 (BHLHE40). *BANCR*, *OSMR*, and *SERPING1* exhibit correlated expression in human embryos (**Fig. 2G**), while *BHLHE40* is upregulated in naïve PSCs compared to primed PSCs (Bayerl et al., 2021). These 26 genes were consistently induced by G-CSF + tamoxifen and LIF + tamoxifen in hESCs, but showed little to no activation when either molecule was used alone (**Fig. 2H**). Notably, expression of classical primed and naïve pluripotency markers (*POU5F1*, *SOX2*, *NANOG*, *ACVR2A*, *ACVR2B*, *SUSD2*, *ESRRB*, *CEACAM1*, *A4GALT*) remained largely unchanged (**Fig. S2C**), indicating that these 26 genes function as early response genes driving the primed-to-naïve transition.

### G-CSF-induced activation of early response genes is dependent on recruitment of JAK kinases to GP130

To elucidate the molecular mechanism underlying G-CSF-induced early response gene activation, we investigated the key intracellular factors recruited to the GCSFR:gp130 chimeric receptor. GP130 signaling is regulated by several well-characterized factors in mouse ESCs, including JAK kinases (Ernst *et al*, 1996), STAT3 (Boeuf *et al*, 1997; Niwa *et al*., 1998), SHP2 phosphatase (Burdon *et al*., 1999), and the YES and HCK kinases (Anneren *et al*, 2004; Ernst *et al*, 1994; Ernst *et al*., 1996). Expression analysis confirmed that all these factors were present in OS3 hESCs (**Fig. 3A**). To dissect their individual roles, we engineered mutant GCSFR:gp130 receptors carrying targeted deletions or point mutations to disrupt specific recruitment sites (**Fig. 3B**). The following receptor variants were used: GCSFR:gp130(Y126-275F), impaired STAT3 recruitment (Niwa *et al*., 1998); GCSFR:gp130(Y118F), disrupted SHP2 binding (Burdon *et al*., 1999); GCSFR:gp130Δ122-171, blocked HCK recruitment (Ward *et al*, 1998); GCSFR:gp130Δ172-187, prevented YES recruitment (Taniguchi *et al*, 2015); GCSFR:gp130Δ13-18, inhibited JAK binding (this study); GCSFR:gp130Δ50-278, deleted the entire intracellular domain except for the JAK-binding site (this paper); GCSFR:gp130Δ13-18/Δ50-278, abolished the recruitment of all signaling factors, including JAK (this study). These receptor mutants were validated via Western blot analysis in 293T cells, confirming their expected phosphorylation patterns for STAT3 and SHP2 (**Fig. S4A**). Next, we generated OS3-derived stable cell lines expressing each mutant receptor under FGF2/KOSR conditions, resulting in the following lines: OS3-GRgp130ΔSHP2, OS3-GRgp130ΔSTAT3, OS3-GRgp130ΔHCK, OS3-GRgp130ΔYES, OS3-GRgp130short, OS3-GRgp130ΔJAK and OS3-GRgp130shortΔJAK.

**Figure 3:**
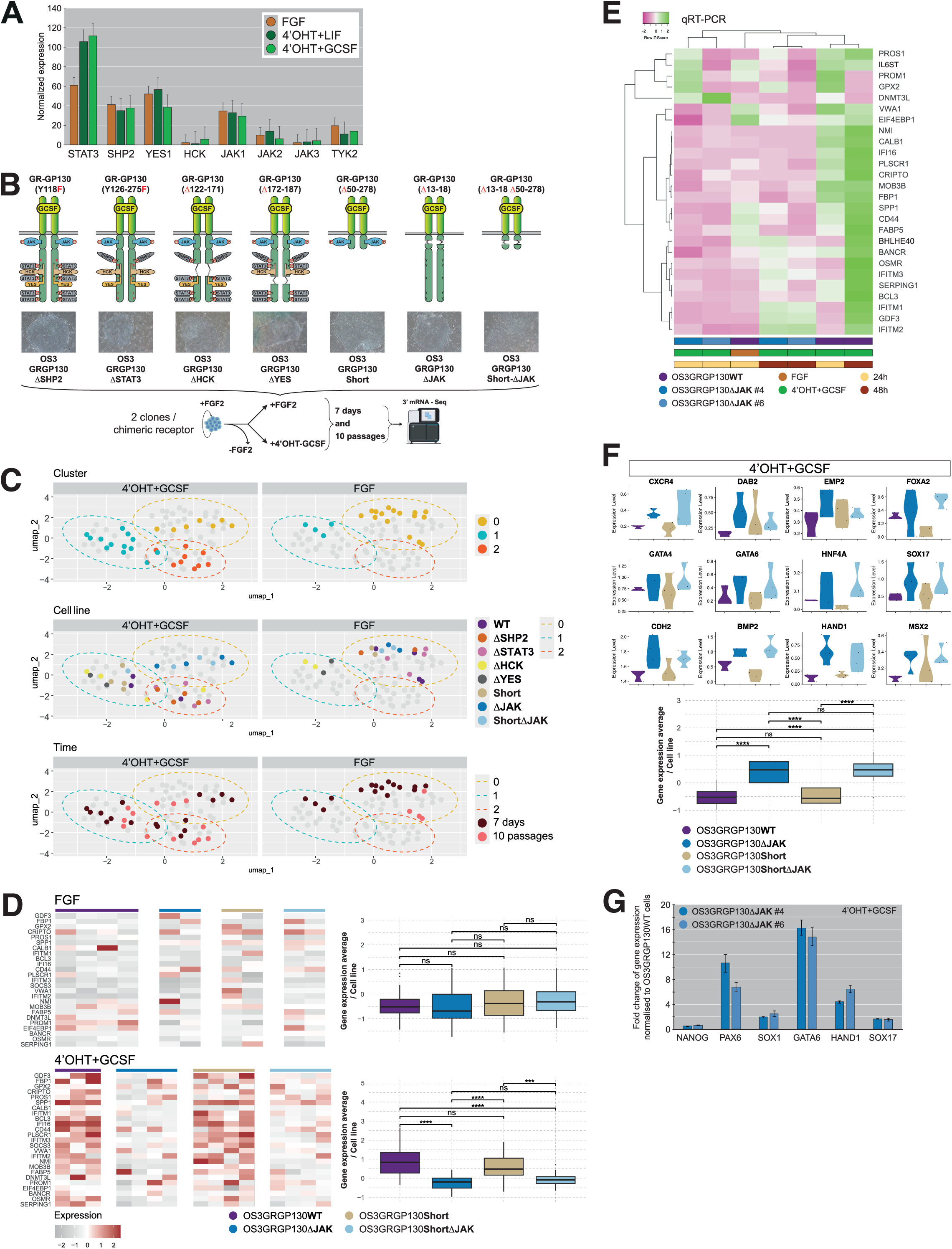
Transcriptomic analysis of mutant OS3-GRgp130 cell lines under G-CSF and tamoxifen treatment. **(A)** Histogram representation of the mRNA level (DESeq2-normalized counts) of STAT3, SHP2, YES, HCK, JAK1/2/3 in OS3-GRgp130wt cells cultured with FGF, 4’OHT+LIF and 4’OHT+GCSF. (**B**) Schematic representation of the seven GCSF:GP130 mutant receptors designed to disrupt specific signaling pathways. Mutant GCSF:GP130(Y126-275F) blocks STAT3 recruitment; GCSF:GP130(Y118F) prevents SHP2 binding; GCSF:GP130(Δ122-171) blocks HCK recruitment; GCSF:GP130(Δ172-187) disrupts YES binding; GCSF:GP130(Δ13-18) inhibits JAK recruitment; GCSF:GP130(Δ50-279) removes the binding sites for STAT3, SHP2, YES, and HCK, while retaining JAK recruitment; and GCSF:GP130(Δ13-18/Δ50-279) eliminates recruitment of all signaling factors, including JAKs. Each mutant receptor was introduced in OS3 cells, resulting in OS3-GRgp130ΔSTAT3, OS3-GRgp130ΔSHP2, OS3-GRgp130ΔSTAT3ΔSHP2, OS3-GRgp130ΔHCK, OS3-GRgp130ΔYES, OS3-GRgp130ΔJAK, OS3-GRgp130short, and OS3-GRgp130shortΔJAK, respectively. Two clones of each cell lines were treated for 7 days and 10 passages with FGF and ‘4OHT+GCSF, followed by RNA sequencing. (**C**) UMAP plot from RNA-seq analysis of the samples described in (A). Top panel: Samples are color-coded by Seurat cluster. Middle panel: Samples are color-coded based on GCSF:GP130 mutant cell lines. Lower panel: Samples are color-coded according to the culture duration. (**D**) Heatmap and quantification boxplot showing the expression of the 25 tamoxifen/G-CSF-induced genes previously identified, in OS3-GRgp130WT, OS3-GRgp130ΔJAK, OS3-GRgp130short, and OS3-GRgp130shortΔJAK cultured in FGF and 4’OHT+GCSF. Expression levels are row-wise z-transformed DESeq2-normalized counts. (**E**) Hierarchical clustering and heatmap of qRT-PCR expression analysis of the 25 tamoxifen/G-CSF-induced genes in OS3-GRgp130WT and OS3-GRgp130ΔJAK (clones #4 and #6) cultured in FGF and 4’OHT+GCSF for 24h and 48h, using Euclidean coefficient as a measure of distance between rows and columns and the complete linkage as a clustering method. (**F**) Violin plot and quantification boxplot showing the expression (DESeq2-normalized counts) of differentiation markers (CXCR4, DAB2, EMP2, FOXA2, GATA4, GATA6, HNF4A, SOX17, CDH2, BMP2, HAND1, MSX2) in OS3-GRgp130WT, OS3-GRgp130ΔJAK, OS3-GRgp130short, and OS3-GRgp130shortΔJAK cultured in 4’OHT+GCSF. (**G**) Histogram representation of the mRNA level (ΔCt) of NANOG, PAX6, SOX1, GATA6, HAND1, SOX17 in OS3-GRgp130WT and OS3-GRgp130ΔJAK (clones #4 and #6) cultured in 4’OHT+GCSF, after normalization to OS3-GRgp130WT (ΔCt=1). Statistical test: t-test (ns = *p* > 0.05, * = *p* ≤ 0.05, ** = *p* ≤ 0.01, *** = *p* ≤ 0.001, **** = *p* ≤ 0.0001).

To determine whether these mutant receptors retained G-CSF responsiveness, we cultured the OS3-derived cell lines without FGF2 and treated them with tamoxifen + G-CSF for (*i*) seven days and (*ii*) 10 passages, followed by transcriptome analysis (**Fig. 3B**). UMAP analysis of 48 samples identified three major transcriptomic clusters: Cluster 0 comprised FGF2-treated samples and a subset of tamoxifen + G-CSF; Cluster 1 predominantly contained tamoxifen + G-CSF; Cluster 2 was exclusively composed of tamoxifen + G-CSF (**Fig. 3C**). After tamoxifen + G-CSF stimulation, most mutant cell lines (OS3-GRgp130wt, OS3-GRgp130ΔSTAT3, OS3-GRgp130ΔSHP2, OS3-GRgp130ΔHCK, OS3-GRgp130ΔYES, and OS3-GRgp130short) clustered together in Clusters 1 and 2, indicating successful transcriptomic reconfiguration. In contrast, OS3-GRgp130ΔJAK and OS3-GRgp130shortΔJAK clustered with FGF2-treated samples (Cluster 0), suggesting that they failed to undergo transcriptomic changes in response to G-CSF. Notably, these transcriptomic patterns were stable even after 10 passages, reinforcing the requirement of JAK recruitment for G-CSF-mediated transcriptomic reprogramming. The interaction between JAK and the GRgp130short mutant was confirmed by Western blot analysis in OS3-GRgp130short cells treated with GCSF (**Fig. S4B**).

To further validate this finding, we examined the expression of 26 previously identified tamoxifen/G-CSF-induced genes. Under FGF2 culture conditions, these genes were expressed at low levels across all cell lines. However, upon tamoxifen + G-CSF treatment, their expression significantly increased in OS3-GRgp130wt and OS3-GRgp130short cells but remained low in OS3-GRgp130ΔJAK and OS3-GRgp130shortΔJAK cells (**Figs. 3D, 3E, S4C**). Importantly, this gene activation was observed after just 7 days, persisted for 10 passages, and was rapidly induced upon FGF2 withdrawal (**Figs. S4D, S4E**). Notably, OS3-GRgp130ΔJAK and OS3-GRgp130shortΔJAK failed to maintain pluripotency and underwent differentiation, as evidenced by the upregulation of early lineage marker genes, including *CXCR4*, *DAB2*, *EMP2*, *FOXA2*, *GATA4*, *GATA6*, *HNF4A*, *SOX17*, *CDH2*, *BMP2*, *HAND1*, and *MSX2* (**Figs. 3F, 3G**).

These results demonstrate that JAK kinase recruitment to the GCSFR:gp130 receptor is essential for the transcriptional activation of tamoxifen/G-CSF-induced early response genes. In contrast, the recruitment of STAT3, SHP2, HCK, and YES is dispensable for this process.

### Regulation of DNA and histone methylation by JAKs

To investigate how JAK kinases regulate early response gene expression, we examined their impact on DNA and histone modifications. In mice, Jak2 has been shown to regulate pluripotency independently of other gp130-associated factors, primarily by reducing 5mC levels through TET enzymes activation and decreasing repressive histone marks (H3K9me3 and H3K27me3) via JMJD2 activation (Wulansari *et al*, 2021). To assess these regulatory effects in our system, we compared 5mC, 5hmC, and H3K9me3 levels in OS3-GRgp130wt and OS3-GRgp130ΔJAK cells three days after switching from FGF2 to G-CSF + tamoxifen. To exclude spontaneously differentiating cells, we restricted the analysis to NANOG-expressing cells. In OS3-GRgp130ΔJAK cells, we observed a significant increase in 5mC levels and a marked decrease in 5hmC levels compared to OS3-GRgp130wt, indicating a disruption of DNA demethylation when JAK recruitment to the GRgp130 receptor was impaired (**Figs. 4A, 4B**). Similarly, H3K9me3 levels were significantly elevated in OS3-GRgp130ΔJAK cells following G-CSF + tamoxifen treatment, suggesting an increase in repressive chromatin modifications in the absence of JAK signaling (**Fig. 4C**). Consistent results were obtained using two different JAK inhibitors, AG490 and SD1029, applied to OS3-GRgp130wt cells during the transition from FGF2 to G-CSF + tamoxifen (**Figs. S5A, S5B, S5C**). To further evaluate the role of JAK in regulating DNA and histone methylation, we compared 5mC, 5hmC, and H3K9me3 levels in OS3-GRgp130wt and OS3-GRgp130short cells under primed conditions and during G-CSF plus tamoxifen–induced reprogramming. In OS3-GRgp130wt cells, G-CSF plus tamoxifen treatment resulted in decreased 5mC levels and increased 5hmC and H3K9me3 levels compared with FGF2 conditions. Notably, OS3-GRgp130short cells exhibited similar changes in 5mC, 5hmC, and H3K9me3 levels (**Figs. 4D, 4E, 4F**). Together these findings highlight a critical role of JAK kinases in regulating DNA methylation and histone modifications, underscoring their importance in early gene activation in our system.

**Figure 4:**
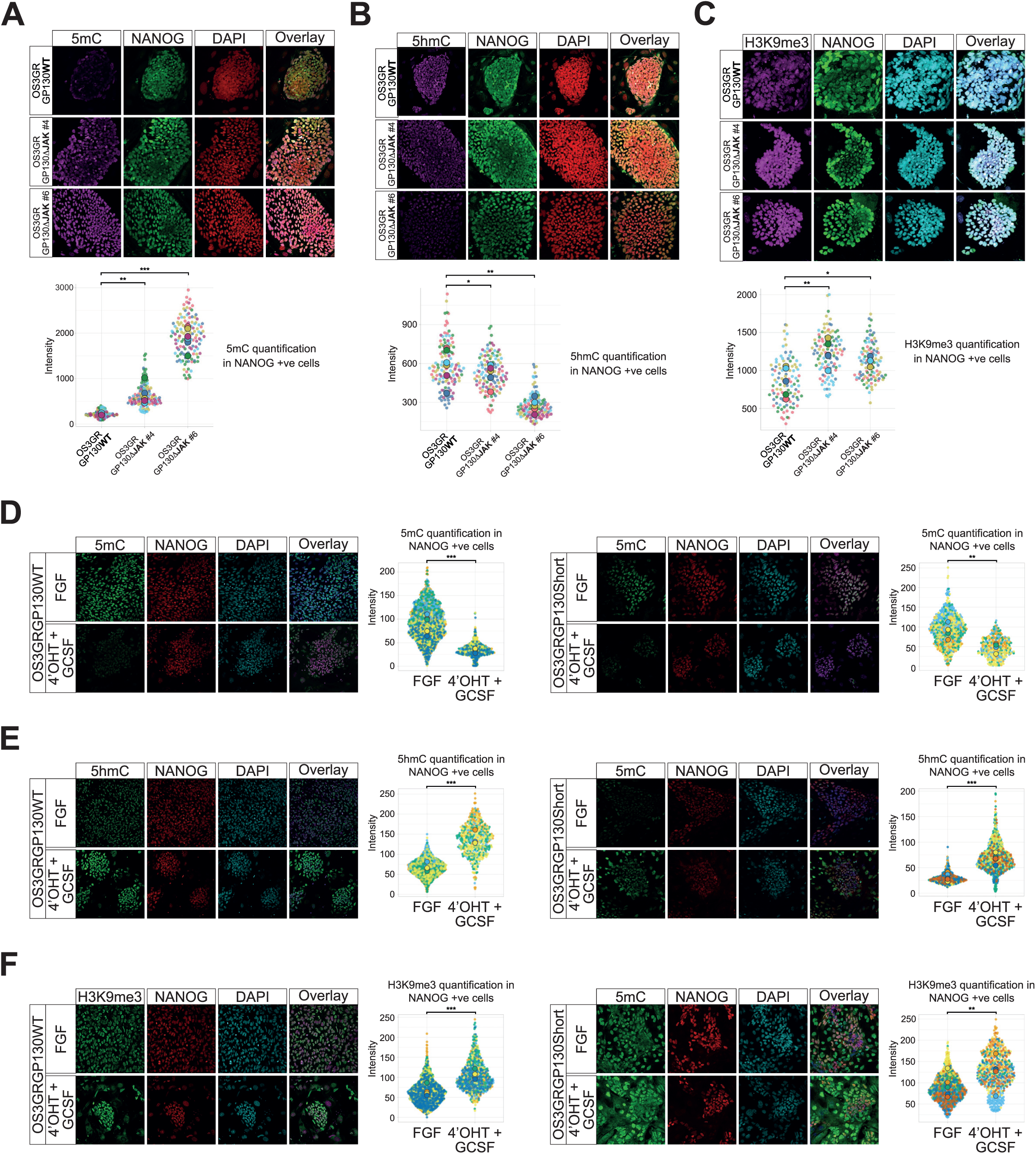
JAK kinases regulate DNA and histone methylation in OS3-GRgp130WT upon G-CSF + tamoxifen stimulation. Immunofluorescence labelling of OS3-GRgp130WT and OS3-GRgp130ΔJAK (clones #4 and #6) cells cultured with 4’OHT+GCSF, with antibodies to NANOG, (**A**) 5mC, (**B**) 5hmC and (**C**) H3K9me3. Immunofluorescence labelling of OS3-GRgp130WT and OS3-GRgp130Short cells cultured either with FGF2 or 4’OHT+GCSF, with antibodies to NANOG, (**D**) 5mC, (**E**) 5hmC and (**F**) H3K9me3.The corresponding histograms show the fluorescence quantification in NANOG positive cells. Statistical test: Welch’s *t*-test (* = *p* ≤ 0.05, ** = *p* ≤ 0.01, *** = *p* ≤ 0.001).

### Effects of early response gene knockdown on self-renewal and pluripotency

To investigate the functional role of early response genes in naive pluripotency reprogramming, we generated stable knockdown OS3-GRgp130wt cell lines under FGF culture conditions using lentiviral vectors carrying validated shRNA sequences targeting individual candidate genes. Following puromycin selection, surviving cells were expanded and subsequently plated at clonal density for seven-day reprogramming with G-CSF and tamoxifen (**Fig. 5A**). Cell morphology was monitored throughout reprogramming for signs of naive-like transition or differentiation. After seven days, pluripotency was assessed via alkaline phosphatase (AP) activity staining and qRT-PCR analysis of pluripotency and lineage-specific gene expression. Among the tested genes, *BCL3*, *CALB1*, *DNMT3L*, *FBP1*, *GDF3*, *GPX2*, *MOB3B*, and *OSMR* showed no visible effects upon knockdown. Reprogrammed cells retained normal morphology, AP staining patterns remained unchanged, and RT-qPCR revealed no significant alterations in lineage-specific gene expression (**Figs. 5B, S6A**). These findings suggest that knockdown of these genes does not impair pluripotency during naive reprogramming. In contrast, knockdown of *BANCR*, *BHLHE40*, *CD44*, *FABP5*, *IFI16*, *IFITM1*, *IFITM2*, *IFITM3*, *IL6ST*, *NMI1*, *PLSCR1, PROM1, PROS1*, and *SERPING1*, *SENP6 / SPP1*, *TDGF1,* and *VWA1* resulted in increased differentiation, as indicated by morphological changes and elevated lineage-specific marker expression (**Figs. 5C, S6B**). Notably, *BHLHE40*, *PROM1*, and *VWA1* knockdown also led to a significant reduction in colony size, suggesting roles in cell proliferation and survival. These results indicate that while some early response genes may be dispensable for naïve reprogramming, others play crucial roles in maintaining self-renewal, proliferation, or suppression of differentiation during the transition to a naïve-like state.

**Figure 5:**
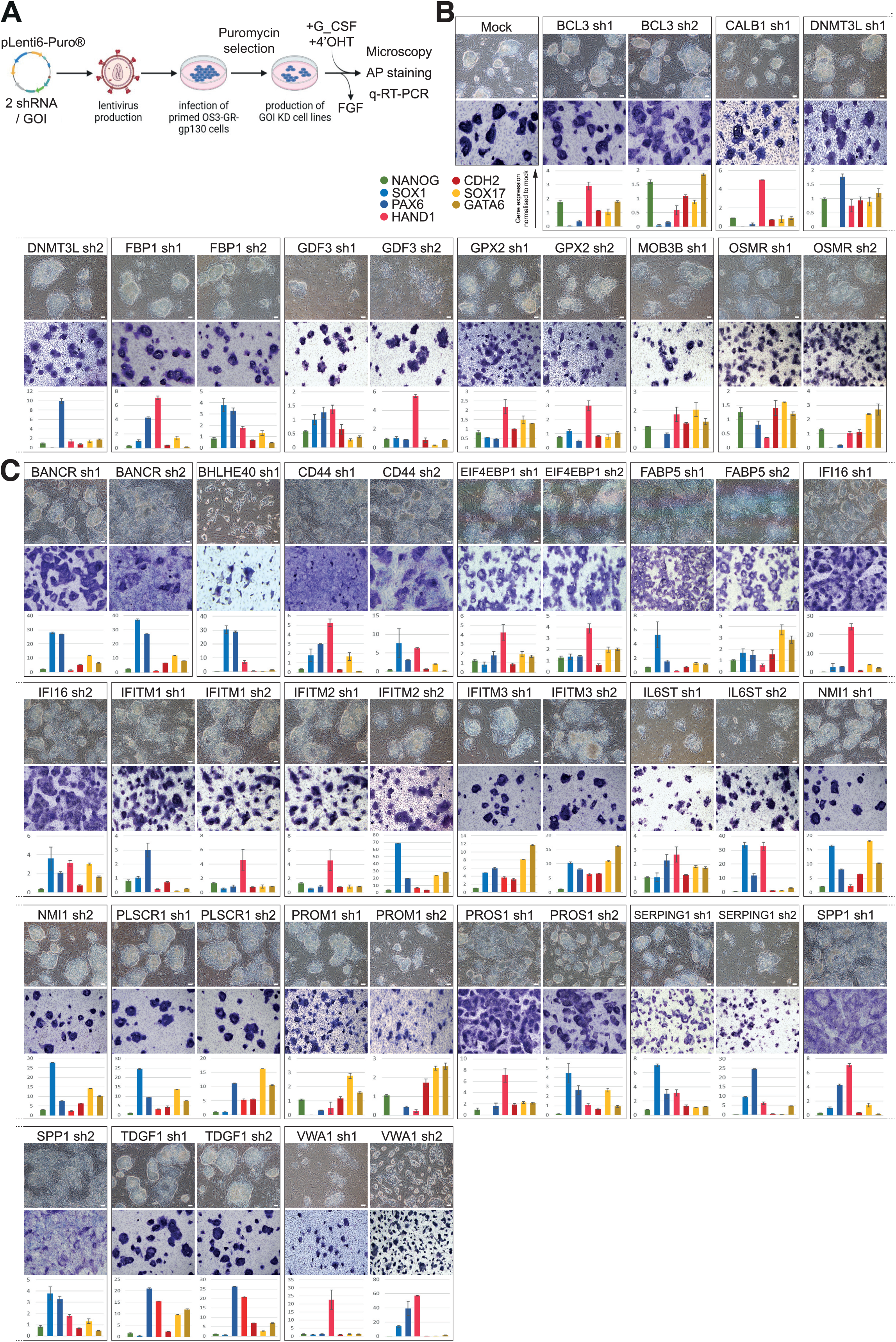
Analysis of candidate gene function in naïve reprogramming. **(A)** Schematic representation of the experimental strategy used to assess the role of each candidate gene in pluripotency. The pLenti6 plasmid contains shRNA sequence under the control of the U6 promoter and a puromycin resistance cassette. Lentiviral particles carrying the two most effective shRNA sequences (Sup. Table1) were used to infect OS3-GRgp130wt cells cultured under FGF conditions. After infection, puromycin selection was applied to eliminate non-transduced cells, and surviving cells were expanded to establish stable knockdown cell lines. To investigate the role of candidate genes in naive reprogramming, these stable knockdown cell lines were treated with G-CSF and tamoxifen for seven days to induce reprogramming into the naïve pluripotent state. GOI: gene of interest. (**B, C**) Phase-contrast microphotographs, AP activity assay, and qRT-PCR analysis of NANOG, SOX1, PAX6, HAND1, CDH2, SOX17 and GATA6 expression in candidate gene knockdown OS3-GRgp130 cell lines cultured with 4’OHT+GCSF for 7 days.

### IFI16 and IFITM1–3 are novel candidate regulators of naïve pluripotency in human PSCs

Four members of the interferon-stimulated gene (ISG) family—IFI16 and IFITM1, IFITM2, and IFITM3—induced substantial differentiation upon knockdown. Notably, these genes are enriched in the human epiblast compared with the primitive endoderm and earlier stages of human embryonic development (**Fig. S7A**). To determine whether they are expressed under established naïve culture conditions, we analyzed published single-cell RNA sequencing (scRNA-seq) datasets from human PSCs reprogrammed in TL2i, PXGL, 5iLA, and 4CL media (Chen *et al.,* 2015, Guo *et al.,* 2021; Moya-Jódar *et al*., 2023; Mazid *et al.,* 2022) (**Fig. S7B**). All four genes were enriched in TL2i- and 4CL-reprogrammed cells. IFITM1, IFITM2, and IFITM3 were enriched in PXGL-reprogrammed cells, whereas IFI16, IFITM2, and IFITM3 were enriched in 5iLA-reprogrammed cells. Together, these observations support a potential role for IFI16 and IFITM1–3 in naïve pluripotency in human PSCs.

To functionally validate the role of interferon-inducible proteins (IFI16, IFITM1, IFITM2, and IFITM3) in pluripotency regulation, we employed a CRISPR-based transcriptional repression strategy (Gil *et al.,* 2024). Human OS3 PSCs (Chen *et al.,* 2015) were engineered to express a doxycycline-inducible dCas9 protein fused to DNMT3A, DNMT3L, and the KRAB repressive domain (**Fig. S8A**). The resulting cell line was designated OS3-CrOff. OS3-CrOff cells were then transfected to stably express three guide RNAs targeting IFI16, IFITM1, IFITM2, or IFITM3. Following hygromycin selection and doxycycline treatment, one guide RNA per gene was validated by qRT-PCR (**Fig. S8B**). The resulting lines were designated OS3-CrOff-IFI16, OS3-CrOff-IFITM1, OS3-CrOff-IFITM2, and OS3-CrOff-IFITM3. To investigate the role of IFI16, IFITM1, IFITM2, and IFITM3 during reprogramming initiation, OS3-CrOff and derivative cell lines were plated at clonal density in FGF2-containing medium. FGF2 was then replaced with LIF and tamoxifen, and cells were cultured for seven days under reprogramming conditions in the presence or absence of doxycycline (**Figs. 6A, S9A**). Colony number and colony size were quantified throughout reprogramming. After seven days, pluripotency was assessed by alkaline phosphatase (AP) staining and qRT-PCR analysis of pluripotency-associated and lineage-specific gene expression. Repression of IFI16 and IFITM3 significantly reduced colony number (**Figs. 6B, S9B**), whereas repression of IFI16, IFITM1, and IFITM3 decreased colony size (**Fig. 6C**). AP staining and qRT-PCR analyses did not reveal induction of differentiation upon repression of IFI16, IFITM1, IFITM2, or IFITM3 (**Figs. 6A, S9C**). These findings indicate that IFI16, IFITM1, and IFITM3 contribute to cell proliferation during early reprogramming without altering the pluripotency–differentiation balance.

**Figure 6.**
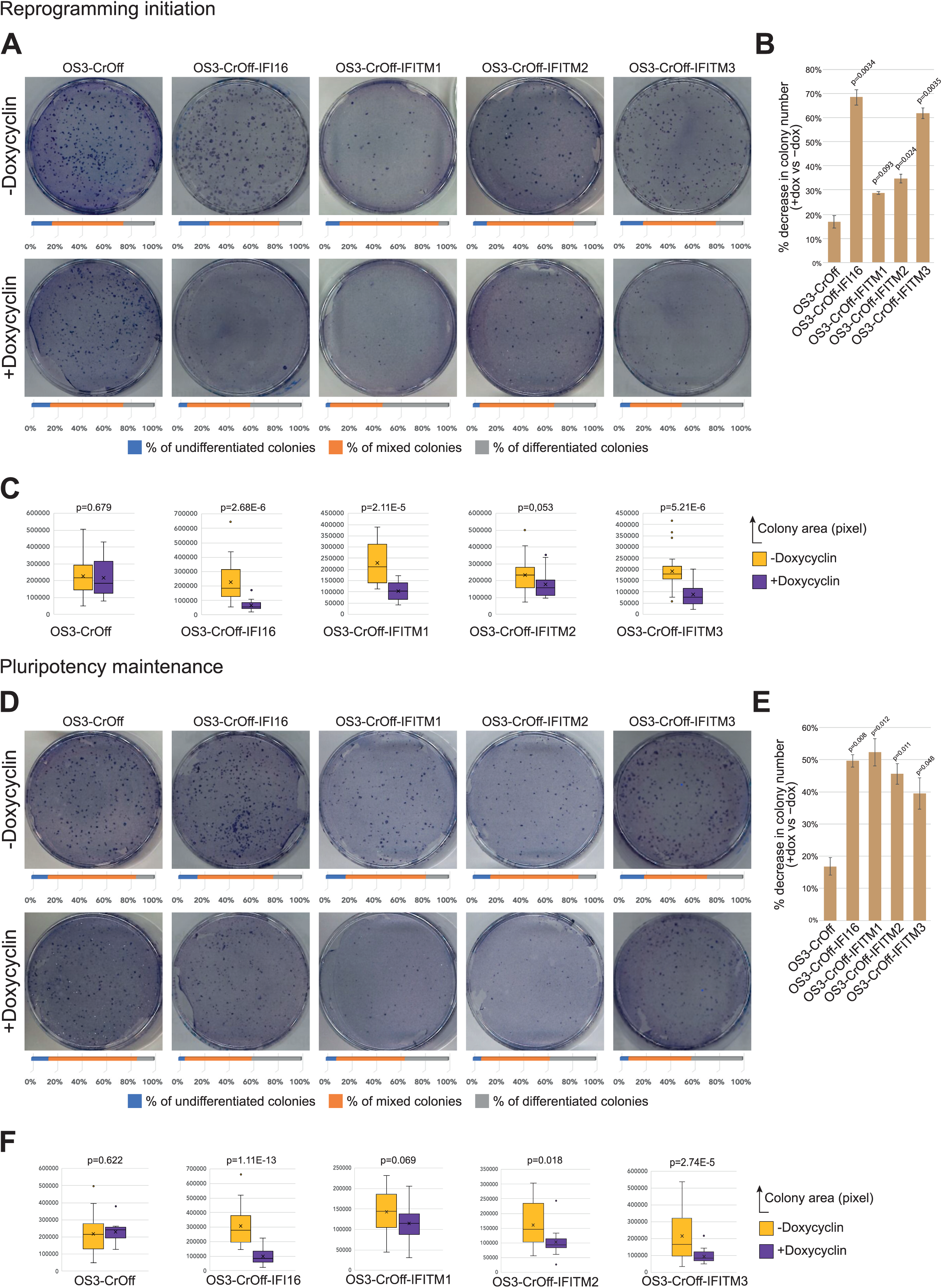
IFI16 and IFITM1–3 regulate proliferation during the establishment of naïve pluripotency. (**A**) Alkaline phosphatase (AP) activity assay performed in OS3-CrOff, OS3-CrOff-IFI16, OS3-CrOff-IFITM1, OS3-CrOff-IFITM2, and OS3-CrOff-IFITM3 cells plated at clonal density in FGF2-containing medium. FGF2 was subsequently replaced with LIF and tamoxifen, and cells were cultured for 7 days in the presence or absence of doxycycline. Histogram bars below each representative image indicate the percentage of undifferentiated, mixed, and differentiated colonies. (**B**) Histogram showing the percentage decrease in colony number between doxycycline-treated (+Dox) and untreated (–Dox) conditions, as quantified in (A). Data represent mean ± SE from two independent replicates. Statistical analysis was performed using Student’s *t*-test. (**C**) Box plot showing colony surface area measured in (A). Statistical analysis was performed using Welch’s *t*-test. (**D**) Alkaline phosphatase (AP) activity assay performed in OS3-CrOff, OS3-CrOff-IFI16, OS3-CrOff-IFITM1, OS3-CrOff-IFITM2, and OS3-CrOff-IFITM3 cells plated at clonal density in Tamoxifen + LIF-containing medium, cultured for seven days in the presence or absence of doxycycline. Histogram bars below each representative image indicate the percentage of undifferentiated, mixed, and differentiated colonies. (**E**) Histogram showing the percentage decrease in colony number between doxycycline-treated (+Dox) and untreated (–Dox) conditions, as quantified in (D). Data represent mean ± SE from two independent replicates. Statistical analysis was performed using Student’s *t*-test. (**F**) Box plot showing colony surface area measured in (D). Statistical analysis was performed using Welch’s *t*-test.

We next assessed the role of IFI16, IFITM1, IFITM2, and IFITM3 in pluripotency maintenance. OS3-CrOff and derivative cell lines were deprived of FGF2 and reprogrammed in LIF- and tamoxifen-containing medium. After five passages, cells were plated at clonal density in LIF and tamoxifen and cultured for an additional seven days in the presence or absence of doxycycline (**Figs. 6D, S9D**). Clonogenicity and pluripotency were evaluated as described above. In contrast to the initiation phase, repression of all four genes during pluripotency maintenance reduced both colony number and colony size (**Figs. 6E, 6F, S9E**), without inducting differentiation (**Figs. 6D, S9F**). These results indicate that all four genes contribute to the regulation of cell proliferation during pluripotency maintenance. Notably, IFITM2 appears to play a more prominent role in the maintenance phase than during reprogramming initiation.

## Discussion

In this study, we identified approximately 50 genes whose expression is upregulated within 24 hours following dual activation of STAT3-ER^T2^ by tamoxifen and JAK kinases via GP130 signaling in primed hESCs. This mode of activation highlights a synergistic interaction between these two pathways. This observation aligns with our previous findings–and is further supported by the present study–that such synergy is required for exit from the primed state of pluripotency and for irreversible commitment to naive pluripotency (Chen *et al*., 2015). Our work provides a first mechanistic insight into this synergy. We propose a model in which JAK kinase activation, triggered by GP130 dimerization, initiates local chromatin opening by reducing DNA methylation and histone H3K9 methylation. This chromatin remodeling might permit subsequent binding of STAT3-ER^T2^ and activation of its target genes. In this model, recruitment of endogenous STAT3 to the GP130 receptor is dispensable, as its role is fulfilled by STAT3-ERT2. This is consistent with findings in mice, where STAT3-ER^T2^ activation alone is sufficient to maintain self-renewal of naïve ESCs, rendering endogenous STAT3 activator via LIF unnecessary (Matsuda *et al*, 1999). However, unlike the situation in mice, where the chromatin is already accessible as a result of continuous self-renewal in the naïve state, in primed hESC cells STAT3-ER^T2^ alone is insufficient. JAK signaling must be activated in parallel to open chromatin at key loci, thereby enabling STAT3-ER^T2^ binding and transcriptional activation.

Among the 50 genes identified, only six genes (*CYP1B1, ICAM1, MYCN, RGS16, SOCS3,* and *ZFP36*) are known STAT3 target in mouse embryonic stem cells (Bourillot *et al*, 2009; Cartwright *et al*, 2005). Notably, none of the naïve pluripotency genes activated by STAT3 in mice, such as *Klf4, Klf5, Gbx2, Tbx3,* or *Tfcp2l1* (Aksoy *et al*, 2014; Bourillot *et al*., 2009; Martello *et al*, 2013; Niwa *et al*, 2009; Qiu *et al*, 2015; Tai & Ying, 2013) were identified. These genes are activated later during the reprogramming of human PSCs to the naïve state. Similarly, human naïve pluripotency markers, such as *KLF17, SUSD2, ALPPL2, DPPA3, DPPA5, CD75,* and *CD130* are not among the genes activated at the onset of naïve reprogramming but are induced later. These observations suggest that the factors required for maintaining naïve pluripotency differ from those that initiate it, the latter being among the genes identified in our study. We specifically focused on 25 genes whose expression strongly correlates with pluripotency in human pre-implantation embryo cells. Among these, knockdown of *BHLHE40, FABP5, PLSCR1, SERPING1, EIF4EBP1, IFI16, IFITM2, IFITM3, IL6ST, PROM1, SPP1,* and *VWA1* expression blocks the initiation of naïve reprogramming in F-OS3-GR-gp130(WT) cells induced by 4’OHT+G-CSF. It would be valuable to test whether these genes also play a role in maintaining naïve pluripotency.

Among the early genes identified and analyzed, several are interferon-stimulated genes (ISGs), including IFITM1, IFITM2, IFITM3, IFI16, PLSCR1, and NMI. These genes encode regulatory elements involved in interferon signaling and perform various cellular functions in normal and cancer cells. Proteins IFITM1, IFITM2, and IFITM3 are induced in response to viral infections, altering membrane properties to block viral entry (Friedlova *et al*, 2022; Gomez-Herranz *et al*, 2023). IFI16, also known as p204 in mice, is a member of the p200 protein family and functions as a sensor of microbial and viral DNA in innate immune responses (Zhao *et al*, 2015). Additionally, PLSCR1 interacts with various viral proteins, either by blocking particle entry or replication (Dal Col *et al*, 2022). The constitutive expression of these proteins in naive pluripotent cells may reflect a natural defense mechanism against pathogens. This defense mechanisms could be particularly relevant for integrative viruses in the context of naive pluripotent cells, which play a critical role in ensuring germline genetic integrity and preserving the species’ genome. Beyond this speculative role, IFITM1, IFITM2, IFITM3, PLSCR1, and IFI16 may have additional functions in pluripotent stem cells. Current knowledge highlights connections between IFITMs and the maintenance and self-renewal of stem cells. For example, Ifitms in mice are expressed in primordial germ cells and regulate their migratory capacity during embryonic development (Tanaka *et al*, 2005). IFITM proteins are also commonly overexpressed in various cancer cells of diverse origins and are associated with most hallmarks of tumor cells such as proliferation, angiogenesis, and metastatic invasion (Friedlova *et al*., 2022; Gomez-Herranz *et al*., 2023). Inactivation of IFITM3 in glioma cells, for instance, induces G0/G1 phase arrest (Zhao *et al*, 2013). IFITM3 also activates STAT3 phosphorylation, promoting tissue invasion by melanoma cells (Wang *et al*, 2020). IFITM1, IFITM2, and IFITM3 can activate PI3K/AKT signaling (Hou *et al*, 2021), a pathway essential for controlling naive pluripotency both *in vitro* (Paling *et al*, 2004; Storm *et al*, 2007) and *in vivo* (Geiselmann *et al*, 2024). Notably, IFITM1 has been shown to repress MAPK signaling and a stabilize p53, two key features of naive PSCs (Yang *et al*, 2007). Similarly, PLSCR1 acts as a p53 activator by interacting with ONZIN to inhibit AKT-mediated MDM2 phosphorylation, leading to p53 stabilization (Li *et al*, 2006). IFI16 also interacts with p53 in human primed PSCs (He *et al*, 2020). Since tight regulation of p53 levels is crucial for maintaining pluripotent cell fitness in the epiblast of the pre-implantation embryo (Bowling *et al*, 2018; Zhang *et al*, 2017), the functional relationships between IFI16, PLSCR1, IFITM1, and p53 represent a promising area for further exploration.

In contrast, little is known about MOB3B. Related genes in the MOB family, such as MOB3A and MOB3C, have been shown to inhibit Hippo/MST/LATS signaling in cancer stem cells. Suppressing MOB3 family gene expression reduces proliferation and tumor growth in cancer cell lines (Dutchak *et al*, 2022). It is plausible that MOB3B functions similarly in human naïve PSCs by regulating Hippo signaling to balance ICM/trophoblast segregation while supporting the proliferation of pluripotent cells.

Prominin-1 (PROM1, also known as CD133) encodes a pentaspan transmembrane protein localized to membrane protrusions. It is well known for its expression in cancer stem cells and various progenitor cells, including those of the bone marrow, liver, kidney, and intestine (Bahn & Ko, 2023). CD133-positive cells derived from colon cancer exhibit long-term expansion in vitro while retaining cancer stem cell (CSC) properties (Ricci-Vitiani *et al*, 2007). PROM1 performs diverse functions in various organs through interactions with multiple molecular partners. For example, PROM1 regulates IL-6-induced liver regeneration via its association with GP130 (Bahn *et al*, 2022). In addition, PROM1 is essential for maintaining CSC properties by activating PI3K and β-Catenin signaling pathways (Mak *et al*, 2012). In naïve hPSCs, PROM1 may similarly enhance GP130 and PI3K activity to support self-renewal.

CD44, a glycoprotein receptor, is functionally associated with CSC renewal in both blood cancers and several types of solid cancers (Sonnentag *et al*, 2024). CD44 activates pathways such as PI3K/AKT, MAPK, and NF-kB, influencing cell migration, proliferation, and survival. Likewise, PROS1, a ligand for TAM receptors, is involved in tissue repair, regeneration, and cancer progression. TAM receptor activation triggers downstream signaling, including PI3K/AKT, which supports cell survival and proliferation (Prieto & Lai, 2024; Wang *et al*, 2021). However, their specific roles in human naïve PSCs remains unclear.

The lncRNA BANCR is primary known for activating the Wnt/β-catenin signaling pathway in cancers such as pancreatic cancer and retinoblastoma through miRNA sponging. BANCR also suppresses p38 MAPK and JNK activation (Hussen *et al*, 2021). Its upregulation during the transition from the primed to the naïve state aligns with the signaling characteristics of naive PSCs.

Finally, OSMR encoding the receptor for Oncostatin M, functions similarly to LIF by inducing heterodimerization with GP130 and recruiting JAK kinases. OSMR can fully substitute for LIF in maintaining naïve pluripotency in mouse ESCs (Ye *et al*, 2021). We propose that OSMR serves as an auxiliary pathway to activate JAK signaling, aiding the transition from the primed to the naïve state in human PSCs.

## Materials and Methods

### Media composition, culture, and electroporation

The male human ES cell line OS3 (Chen *et al*., 2015) and their clonal derivatives were routinely cultured at 37 °C in 5% CO_2_ and 5% O_2_ in knockout Dulbecco’s modified Eagle’s medium (KO-DMEM) supplemented with 20% knockout serum replacement (KOSR), 1 mM glutamine, 0.1 mM β-mercaptoethanol (Sigma), 1% non-essential amino acid (Gibco), and 4–5 ng/mL FGF2 (Gibco) on growth-inactivated murine embryonic fibroblasts. They were passaged by mechanical dissociation using collagenase I. After withdrawal of FGF2 and addition of 10,000 U/ml LIF, 3 µM G-CSF, and 250 nM 4’hydroxy-tamoxifen, cells were enzymatically passaged using 1X trypsin (Gibco), with 10 µM ROCK inhibitor Y27632 (Calbiochem) added for 24 hours after each passage.

To generate PiggyBac (PB) transgenic lines, 10^6 cells were co-transfected using the Neon electroporation system (Invitrogen; 1050 V, 20 ms, 2 pulses) with 2.5 μg of the PB plasmid and 2.5 μg of the PBase-expressing vector *pCAGPBase* (Wang et al., 2008). Stable transfectants were selected with G418 (250 μg/mL) for 7 days. Following clonal selection, individual clones were verified by PCR and maintained under G418 selection (250 μg/mL).

To generate Sleeping Beauty (SB) transposon cell lines, 10^6 cells were co-transfected using the Neon electroporation system (Invitrogen; 1050 V, 20 ms, 2 pulses) with 2.5 μg of the SB plasmid and 2.5 μg of the SB100X transposase-expressing vector *pCMV(CAT)T7-SB100* (Addgene #127909). Stable transfectants were selected with hygromycin (200 μg/mL) for the *pSB-TRE-CRISPRoff-EF1A-TetOn* plasmid (Addgene #203355) and puromycin (1 μg/mL) for the *pSB-BbsI-sgRNA* plasmid (Addgene #203359) for 7 days. Following clonal selection, clones overexpressing CRISPRoff dCas9 were confirmed by western blot and maintained under hygromycin selection (200 μg/mL). OS3-CrOff-sgRNA clones were subsequently verified by qPCR and cultured under dual selection with hygromycin (200 μg/mL) and puromycin (1 μg/mL).

### Plasmid constructs and shRNA design

Numbering begins at the first residue of the 278 amino-acid intracellular domain of mouse gp130.

- The cDNA sequences encoding GCSFR:gp130(WT) and GCSFR:gp130(Y118F) were PCR amplified from plasmids *pPB-CAG-GCSFR:gp130* and *pPB-CAG-GCSFR:gp(Y118F)* (Niwa *et al*., 1998), using primers with *NheI* and *EcoRV* restriction sites. The resulting fragments were subcloned into the *NheI* and *BamHI* (blunt) sites of *pPB-CAG-MCS-IRES-Neo3*, to generate *pPB-CAG-GCSFR:gp130(WT)-IRES-Neo3* and *pPB-CAG-GCSF:gp130(Y118F)-IRES-Neo3*.
- To generate *pPB-CAG-GCSFR:gp130(Y126-275F)-IRES-Neo3* plasmid, a 5’-*NheI/NsiI*-3’ fragment from *pPB-CAG-GCSFR:gp130(WT)-IRES-Neo3* was combined with a synthetic 5’-*NsiI/EcoRV*-3’ DNA fragment (GeneArt) containing four tyrosine-to-phenylalanine substitutions (Y126F, Y173F, Y265F, and Y275F). Both fragments were subcloned between the *NheI* and *BamHI* (blunt) sites of the *pPB-CAG-MCS-IRES-Neo3* vector.
- To generate *pPB-CAG-GCSFR:gp130(Δ122-171)-IRES-Neo3* and *pPB-CAG-GCSFR:gp130(Δ172-187)-IRES-Neo3*, 5’-*BstEII/BsrGI*-3’ fragments containing the Δ122-171 and Δ172-187 deletions were generated using nested PCR and subsequently subcloned into the *BstEII* and *BsrGI* sites of *pPB-CAG-GCSFR:gp130(WT)-IRES-Neo3* plasmid.
- To generate *pPB-CAG-GCSFR:gp130(Δ50-278)-IRES-Neo3,* a fragment encoding the extracellular domain of human GCSFR and the first 50 amino acids of the intracellular domain of mouse gp130 was PCR-amplified from *pPB-CAG-GCSFR:gp130(WT)* with primers containing *NheI* and *EcoRV* sites. This fragment was subcloned into the *NheI* and *BamHI* (blunt) sites of *pPB-CAG-MCS-IRES-Neo3* to generate *pPB-CAG-GCSFR:gp130(Δ50-278)-IRES-Neo3*.
- To generate *pPB-CAG-GCSFR:gp130(Δ13-18)-IRES-Neo3*, a 5’-*SfiI/BsrGI*-3’ fragment comprising the Δ13-18 deletion was generated by nested PCR and subcloned into the *SfiI* and *BsrGI* sites of *pPB-CAG-GCSFR:gp130(WT)-IRES-Neo3*.
- To generate *pPB-CAG-GCSFR:gp130(Δ13-18)(Δ50-278)-IRES-Neo3*, a fragment encoding the extracellular domain of human GCSFR and the first 43 amino acids of the intracellular domain of gp130, containing the Δ13-18 deletion, was PCR-amplified from *pPB-CAG-GCSFR:gp130(Δ13-18)-IRES-Neo3* with primers containing *NheI* and *EcoRV* sites. The resulting fragment was subcloned into the *NheI* and *BamHI* (blunt) sites of *pPB-CAG-MCS-IRES-Neo3* to generate the *pPB-CAG-GCSFR:gp130(Δ13-18)(Δ50-278)-IRES-Neo3*.

For shRNA-mediated knockdown, three independent sequences were designed for each target gene using the BLOCK-iT^TM^ RNAi Designer (ThermoFisher Scientific, [RNAi Designer](https://rnaidesigner.thermofisher.com/rnaiexpress/)). These sequences were cloned into the pENTR^TM^/U6 vector (ThermoFisher), followed by subcloning into the *pLenti6/BLOCK-iT-PGKpuro^R^*vector. The resulting pLenti6-shRNA-puro plasmids were used to produce lentiviral particles and transduce OS3GRgp130 cells reprogrammed with 4’OHT/G-CSF. Real-time PCR was used to measure the level of interference. Details of the shRNA sequences and their interference efficiencies are provided in **Supplementary Table 1**.

For CRISPRoff targeting, three independent sgRNA spacer sequences were designed to target *IFI16*, *IFITM1*, *IFITM2*, and *IFITM3* using the CRISPick web portal (https://portals.broadinstitute.org/gppx/crispick/public). Annealed sense and antisense oligonucleotides containing the appropriate BbsI overhangs were ligated into the BbsI site of the *pSB-BbsI-sgRNA* plasmid. Details of the spacer sequences are shown in **Fig. S8B**.

### Virus production and infection of hESCs

293T cells were transfected with a DNA mixture containing 0.7 µg of the *pMD2.G* plasmid encoding the vesicular stomatitis virus glycoprotein envelope, 1.3 µg of *psPAX2* plasmid encoding the gag, pol, tat, and rev proteins, and 2 µg of the pLenti6-shRNA-puro plasmid carrying the lentiviral genome (Negre *et al*, 2000) using the calcium phosphate precipitation technique. The following day, cells were incubated with 1.2 mL of fresh decomplemented medium and further cultured for 24 h. The supernatant was then collected, filtered (0.45µm), cleared by centrifugation (3000 rpm, 15 min).

Prior to infection, 2 x 10^5^ OS3-GRgp130wt cells were dissociated with Trypsin-EDTA, and cells were seeded in P6 well containing 1 mL of primed culture medium supplemented with ROCK inhibitor (ROCKi) and 1 mL of filtered supernatant containing lentiviral particles. The P6 plate was sealed with parafilm and centrifuged for 45 minutes at 1200 rpm. After centrifugation, the plate was incubated for 70 minutes before the medium was refreshed with primed culture medium supplemented with ROCKi. After 24 hours, the medium was refreshed daily with primed culture medium supplemented with puromycin. Alkaline phosphatase (AP) activity was detected using the AP substrate kit (Sigma-Aldrich), following the manufacturer’s protocol.

### Immunofluorescence analysis of cells

Cells were fixed in 4% PFA for 20 min at room temperature. After three washes in phosphate-buffered saline (PBS), they were permeabilized in PBS-0.5% TritonX100 for 30 min and blocked in 2% BSA for 1 h at room temperature. For 5’methylcytosine immunolabeling, cells were permeabilized in PBS-0.5% Triton for 15 min, washed in PBS for 20 min, then incubated in 2M HCl for 30 min prior blocking as above. In all cases, cells and embryos were subsequently incubated with primary antibodies diluted in blocking solution overnight at 4°C. Primary antibodies include: NANOG (R&D Systems), H3K9me3 (Abcam), 5mC (Diagenode), and 5hmC (Activemotif). After three washes (3 x 15 min) in PBS, cells were exposed to goat anti-mouse, anti-rabbit immunoglobulin G, or immunoglobulin M conjugated either to fluorescein isothiocyanate, Alexa-488, -555, or -647 (dilution 1:1,000) (Invitrogen) for 1 h at room temperature, followed by nuclear staining with DAPI (Invitrogen, diluted 1:5000 in the blocking solution) for 10 minutes or propidium iodide (PI, Calbiochem, diluted 1:1000 in the blocking solution) for 40 minutes at RT. After three washes in PBS, coverslips were mounted on slides. The cells on coverslips were analyzed using a confocal laser scanning system (Confocal SP5, Leica Microsystems). For immunofluorescence analysis, Welch’s unequal variances *t*-test was employed. The SuperPlotsOfData tool (Postma & Goedhart, 2019) was utilized to account for all measurements and biological replicates in these comparisons.

### Western Blotting

Cells were lysed in cold RIPA lysis buffer (0.5% NP-40, 1% Triton-X, 10% glycerol, 20 mM Hepes pH 7.4, 100 mM NaCl, 1 mM sodium orthovanadate, 0.1% DTT, and protease inhibitors (Roche, #05 892 970 001) for four hours at 4°C. Lysates were cleared by centrifugation for 15 min and stored at -80°C. Protein concentrations were measured using Bradford assay. For SDS-PAGE electrophoresis, 30µg of total proteins were loaded onto each well of Mini-PROTEAN TGX Stain-Free Precast Gels (10%, Biorad, # 4568031), and migrated for 45 min at 120 Volts. Precision Plus Protein Dual Color Standards (Biorad, # 1610374) were used as protein ladder. After electroporation, proteins were transferred onto membranes using Trans-Blot^®^ Turbo^TM^ RTA Midi 0.2 µm Nitrocellulose Transfer Kit (Biorad, #1704271). The membranes were subsequently blocked in TBST solution (200 mM Tris-HCl, 1.5 M NaCl, 0.1% Tween-20, 5% milk) for one hour at room temperature, prior to incubation with primary antibodies diluted in TBST at 4°C for 12 hours. Primary antibodies are listed in **Supplementary Table 2**. Membranes were incubated with HRP-conjugated secondary antibody (Jackon ImmunoResearch anti-mouse ref 211-032-171 and anti-goat ref 115-035-146, dilution 1:5000) for one h at room temperature. After serial washing in TBST, HRP activity was revealed using Clarity^TM^ Western ECL substrate (Biorad, #170-5060) and ChemiDoc^TM^ MP imaging system (Biorad).

### RNA extraction and RNA sequencing

Total RNA was isolated using RNeasy mini kit (Qiagen #74106) with a DNase I (Qiagen #79254) treatment. For the early response genes identification, the 3’ seq RNA profiling (SRP) libraries are prepared from 10ng of total RNA per sample according to Soumillon et al.^1^ In brief, template-switching reverse transcription is used to tag mRNA poly(A) tails with universal adapters, well-specific barcodes, and Unique Molecular Identifiers (UMIs). Ninety-six RNA samples are processed in a 96-well plate with well-specific adapters (E3V6NEXT). Reverse transcription utilizes Maxima H Minus Reverse Transcriptase and specific adapters (E5V6NEXT). After pooling and purification, cDNAs are treated with Exonuclease I and amplified by 12-cycle single-primer PCR using SINGV6. Amplified cDNAs are fragmented using a bead-linked transposome (Illumina). Libraries are prepared with Nextera™ DNA Flex, using modified primers and 8 PCR cycles. Library size (400–800 bp) is verified using an Agilent Tape Station. Quantification of P5-P7 adapters is performed via qPCR. Libraries concentration must be over 15 nM for sequencing. Libraries are sequenced on Illumina NovaSeq 6000 with specific read lengths (e.g., 17-8-105 cycles). Demultiplexing was performed using bcl2fastq2 (version 2.18.12) and adapters were trimmed with Cutadapt (version 1.15 and 3.2). Reads were mapped to the human genome GRCh38 using BWA aligner. Unique UMIs per gene/sample are counted from alignment files. Reads are excluded if: UMIs contain ’N’s, alignments contain more than 3 mismatches, sequences align equally to multiple genes. Number of reads ranged from 0.3 to 4.5 million post-counting.

For the transcriptome analysis of the chimeric receptors, 40ng of total RNA were used for amplification using the QUANT^TM^ SEQ^POOL^ kit (LEXOGEN) according to the manufacturer’s recommendations. cDNA quality was assessed on an Agilent Bioanalyzer 2100, using an Agilent High Sensitivity DNA Kit. Libraries were prepared from 0.15 ng cDNA using the Nextera XT Illumina library preparation kit and sequenced (Paired-end 70-22 bp) on an Illumina NextSeq500 instrument, using a NextSeq 500 High Output 75 cycles kit. FASTQ files are generated using bcl2fastq2 (version 2.18.12) and demultiplexing was performed using *idemux* (Lexogen). Adapters were trimmed with Trimmomatic (https://github.com/timflutre/trimmomatic) so that only reads longer than 50 bp were kept. Number of reads ranged from 0.3 to 7.2 million post adapter trimming. Reads were mapped to the human genome hg38 using STAR aligner (https://github.com/alexdobin/STAR).

### Bioinformatics analysis

All analyses were executed using R software (version 4.4.1). Data normalization and gene expression levels were performed using the DESeq2 R package (version 1.44.0) (Love *et al*, 2014). After log2 transformation, normalized counts were used to produce UMAP and heatmaps with the seurat R package (version 5.1.0). DESeq2 computed log2 transformed Fold Change (FC) and p-values for each feature. Differentially expressed genes (DEGs) were determined by Over-Representation Analysis (ORA) using DESeq2 results and the following thresholds: FC > 2 ∪ FC < (−2), p-value < 0.05. Volcano plots were designed using the ggplot2 package (version 3.5.1). Correlation expression values from Meistermann et al. (Ref) were used to produce correlation heatmap with the pheatmap R package (version 1.0.12).

### Data availability

The RNA-Seq datasets will be publicly available as of the date of publication.

## Acknowledgements

This work has benefited from the facilities and expertise of the Genomics Core Facility GenoA, member of Biogenouest and France Genomique and of the Bioinformatics Core Facility BiRD, member of Biogenouest and Institut Français de Bioinformatique (IFB) (ANR-11-INBS-0013). This work was supported by the Fondation pour la Recherche Médicale (EQU202303016295 to P.S.), the LabExs (ANR-10-LABX-73, REVIVE; ANR-10-LABX-0061, DEVweCAN; ANR-11-LABX-0042, CORTEX), and the University of Lyon within the program “Investissements d’Avenir” (ANR-11-IDEX-0007).

## Author contributions

Investigation: T.K., C.S., N.D., A.K., A.B., M.D., C.R., M.W., A.B.

Formal analysis: P.-Y.B, L.D.

Writing original draft, Funding acquisition: P.-Y.B., P.S.

Conceptualization, Supervision, Validation, Visualization, Project administration, Manuscript review and editing: P.-Y.B., P.S., M.W., L.D.

Resources: L.D.

---

**Supplementary figure 1:**
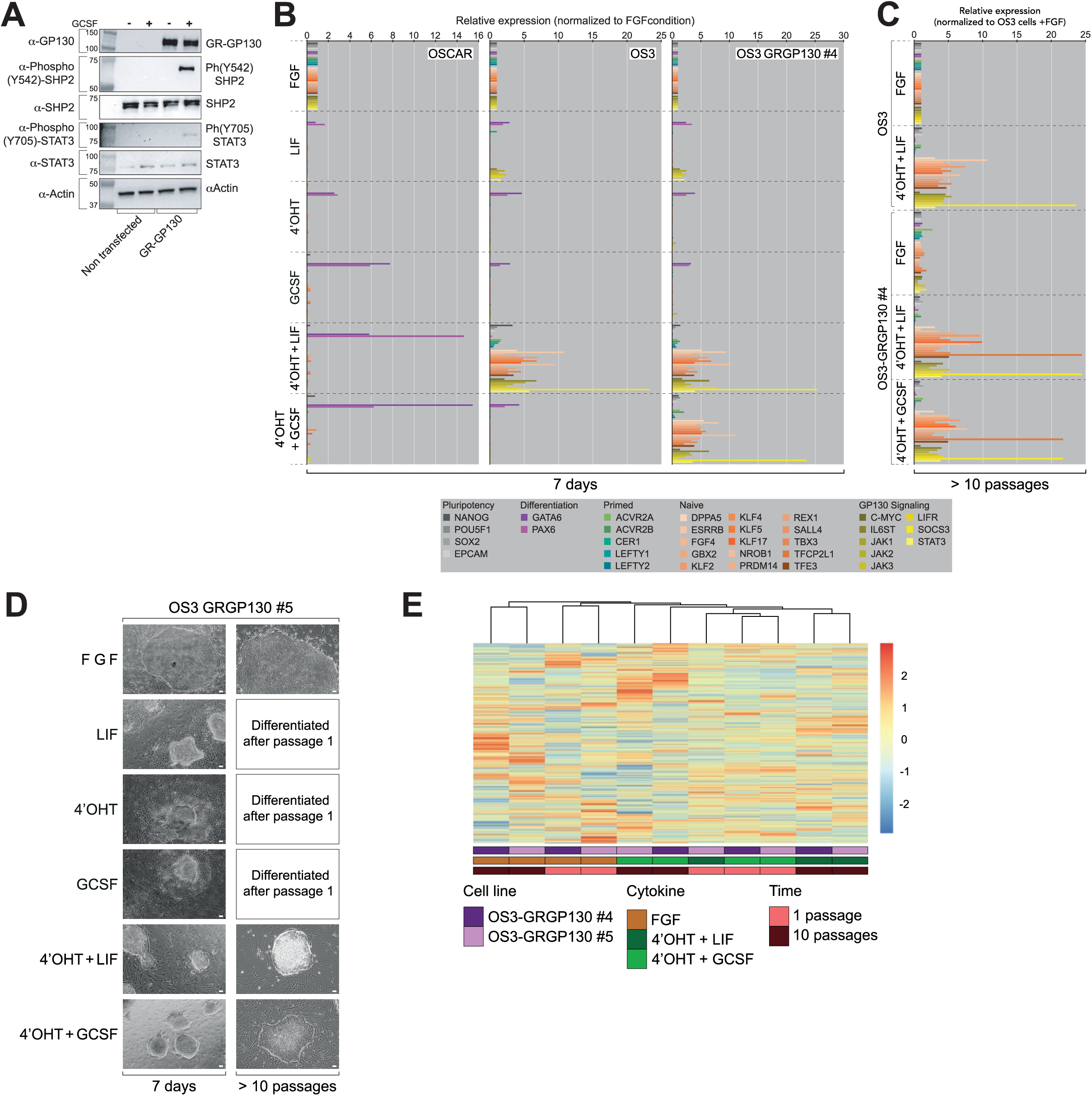
Analysis of gp130/SHP2/STAT3 Signaling and Gene Expression Following G-CSF Stimulation. (**A**) Western blot analysis of GP130, phospho-(Tyr542)-SHP2, SHP2, phospho-(Tyr705)-STAT3, STAT3 expression in HEK 293T cells transfected with the *pPB-CAG-GCSFR:gp130(WT)-IRES-Neo3* plasmid, without stimulation of G-CSF and after stimulation with G-CSF. Histogram representation of the mRNA level (ΔCt) of pluripotency, primed pluripotency, naive pluripotency, differentiation and GP130 signaling markers in OSCAR, OS3 and OS3-GRgp130wt #4 cells treated with FGF, LIF, 4’OHT, GCSF, 4’OHT+LIF or 4’OHT+GCSF for 7 days (**B**), in OS3 and OS3-GRgp130wt #4 cells cultured with FGF, 4’OHT+LIF or 4’OHT+GCSF, over 10 passages (**C**), after normalization to FGF (ΔCt=1). (**D**) Phase-contrast microphotography of OS3-GRgp130wt #5 cells cultured with FGF, LIF, 4’OHT, GCSF, 4’OHT+LIF or 4’OHT+GCSF, for 7 days or over 10 passages. Scale Bar, 50µm. (**E**) Hierarchical clustering and heatmap of transcriptome data (mean values/cell category most differentially expressed 1,000 probe sets) using Pearson correlation coefficient as a measure of distance between rows and Spearman correlation coefficient as a measure of distance between columns.

**Supplementary figure 2:**
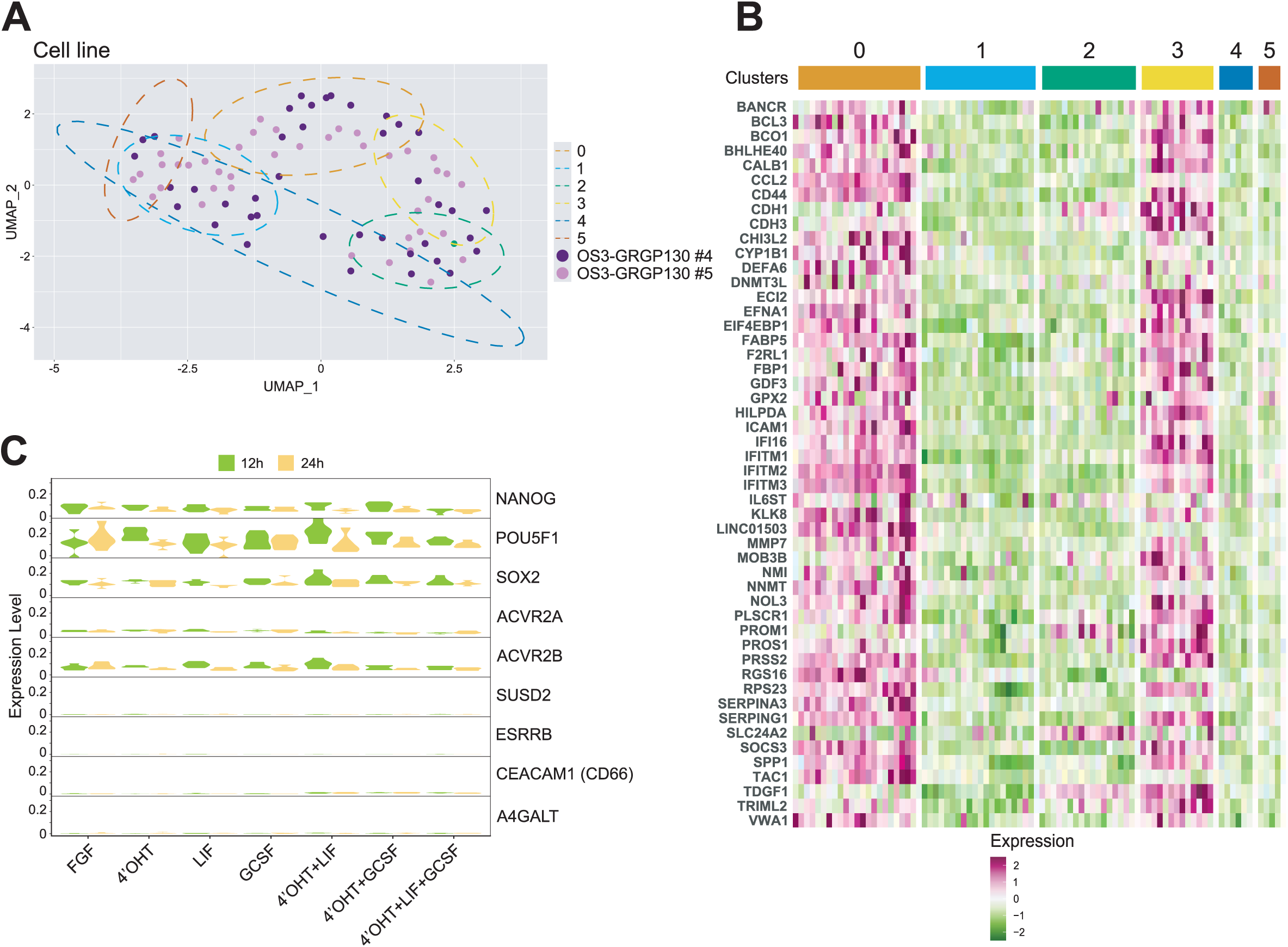
UMAP Clustering and Differential Gene Expression Analysis of OS3-GRgp130wt Cells. (**A**) UMAP plot from RNA-seq analysis of samples described in Fig. 2A. Samples are coloured according to the OS3-GRgp130wt clone. (**B**) Heatmap showing the expression of the 100 most differentially expressed genes identified in **Figs. 2E, S3**. Samples are grouped according to the Seurat clusters. Expression levels are row-wise z-transformed DESeq2-normalized counts. (**C**) Violin plot expression (DESeq2-normalized counts) of NANOG, OCT4, SOX2, ACVR2A, ACVR2B, SUSD2, ESRRB, CEACAM1, A4GALT, in OS3-GRgp130wt cells, across the different culture conditions. Samples are split according the culture time.

**Supplementary figure 3:**
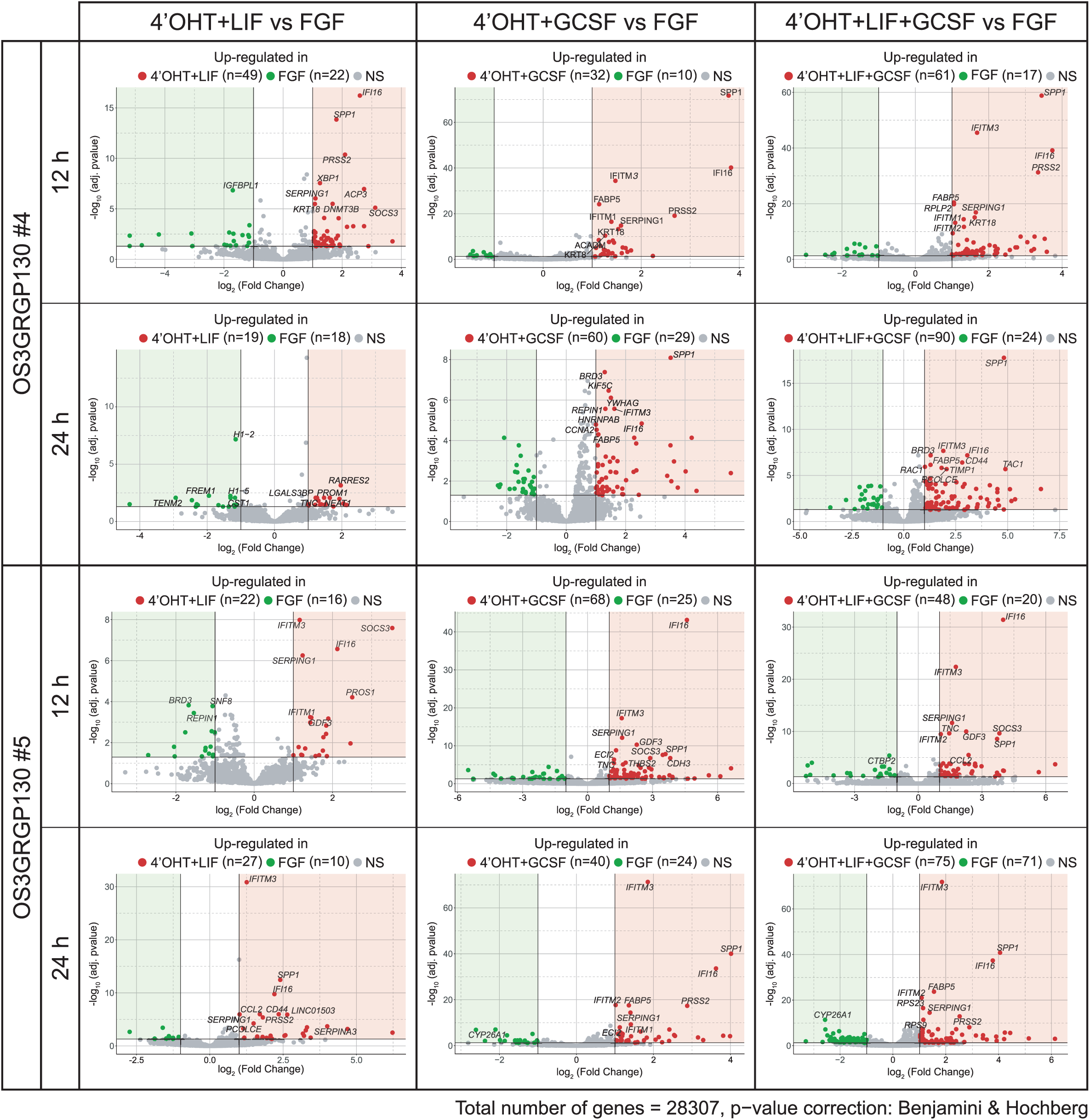
Early Transcriptional Responses to LIF and G-CSF Stimulation in OS3-GRgp130wt Cells. Volcano plot showing the up-regulated genes identified in pairwise comparisons between LIF + 4’OHT, G-CSF + 4’OHT, and LIF + G-CSF + 4’OHT versus FGF2, in OS3-GRgp130wt cells after 12 and 24 hours of treatment. DEG thresholds: FC >2 U FC< (−2), p-value <0.05. For each comparison, the selected genes represent the 10 most differentially expressed ones.

**Supplementary figure 4:**
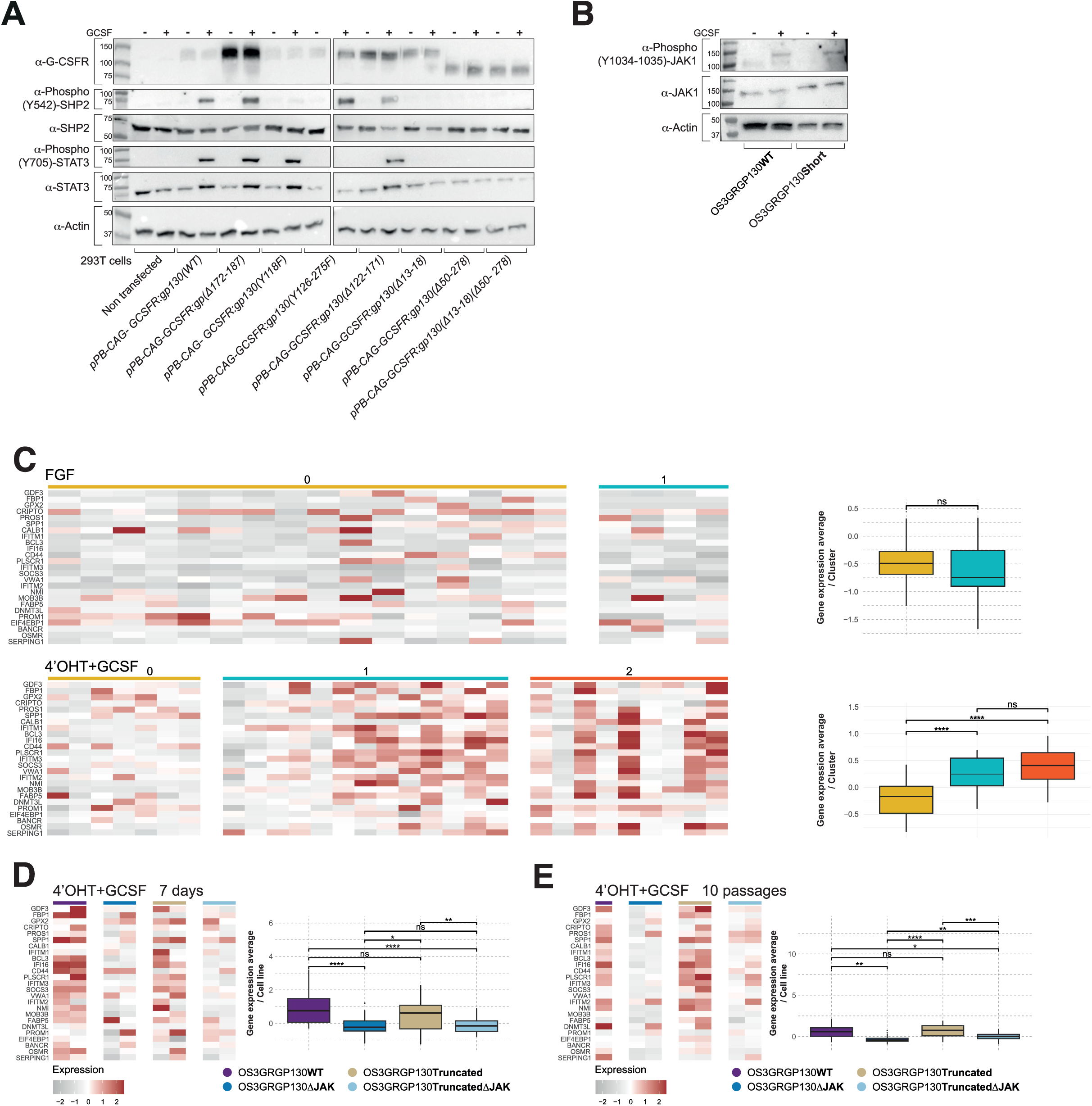
Signaling and Transcriptional Consequences of gp130 Mutations in Response to G-CSF. (**A**) Western blot analysis of GP130, phospho-(Tyr542)-SHP2, SHP2, phospho-(Tyr705)-STAT3, STAT3 expression in HEK 293T cells transfected with the *pPB-CAG-GCSFR:gp130(WT)-IRES-Neo3, pPB-CAG-GCSFR:gp(Δ172-187)-IRES-Neo3*, *pPB-CAG-GCSFR:gp130(Y118F)-IRES-Neo3, pPB-CAG-GCSFR:gp130(Y126-275F)-IRES-Neo3*, *pPB-CAG-GCSFR:gp130(Δ122-171)-IRES-Neo3*, *pPB-CAG-GCSFR:gp130(Δ13-18)-IRES-Neo3, pPB-CAG-GCSFR:gp130(Δ50-278)-IRES-Neo3* and *pPB-CAG-GCSFR: gp130(Δ13-18)(Δ50-278)-IRES-Neo3* plasmids, without stimulation of G-CSF and after stimulation with G-CSF. (**B**) Western blot analysis of phospho-(Y1034-1035)-JAK1 and JAK1 expression in OS3-GRgp130WT and OS3-GRgp130short cells prior to and after stimulation with G-CSF. (**C**) Heatmap and quantification boxplot showing the expression of the 25 tamoxifen/G-CSF-induced genes previously identified, in OS3-GRgp130WT, OS3-GRgp130ΔJAK, OS3-GRgp130short, and OS3-GRgp130shortΔJAK cultured in FGF and 4’OHT+GCSF. Samples are categorized according to Seurat clustering. Expression levels are row-wise *z*-transformed DESeq2-normalized counts. Heatmap and quantification boxplot showing the expression of the 25 tamoxifen/G-CSF-induced genes previously identified, in OS3-GRgp130WT, OS3-GRgp130ΔJAK, OS3-GRgp130short, and OS3-GRgp130shortΔJAK cultured in 4’OHT+GCSF for 7 days (**D**) and 10 passages (**E**). Expression levels are row-wise z-transformed DESeq2-normalized counts. Statistical test: *t*-test (ns = *p* > 0.05, * = *p* ≤ 0.05, ** = *p* ≤ 0.01, *** = *p* ≤ 0.001, **** = *p* ≤ 0.0001.

**Supplementary figure 5:**
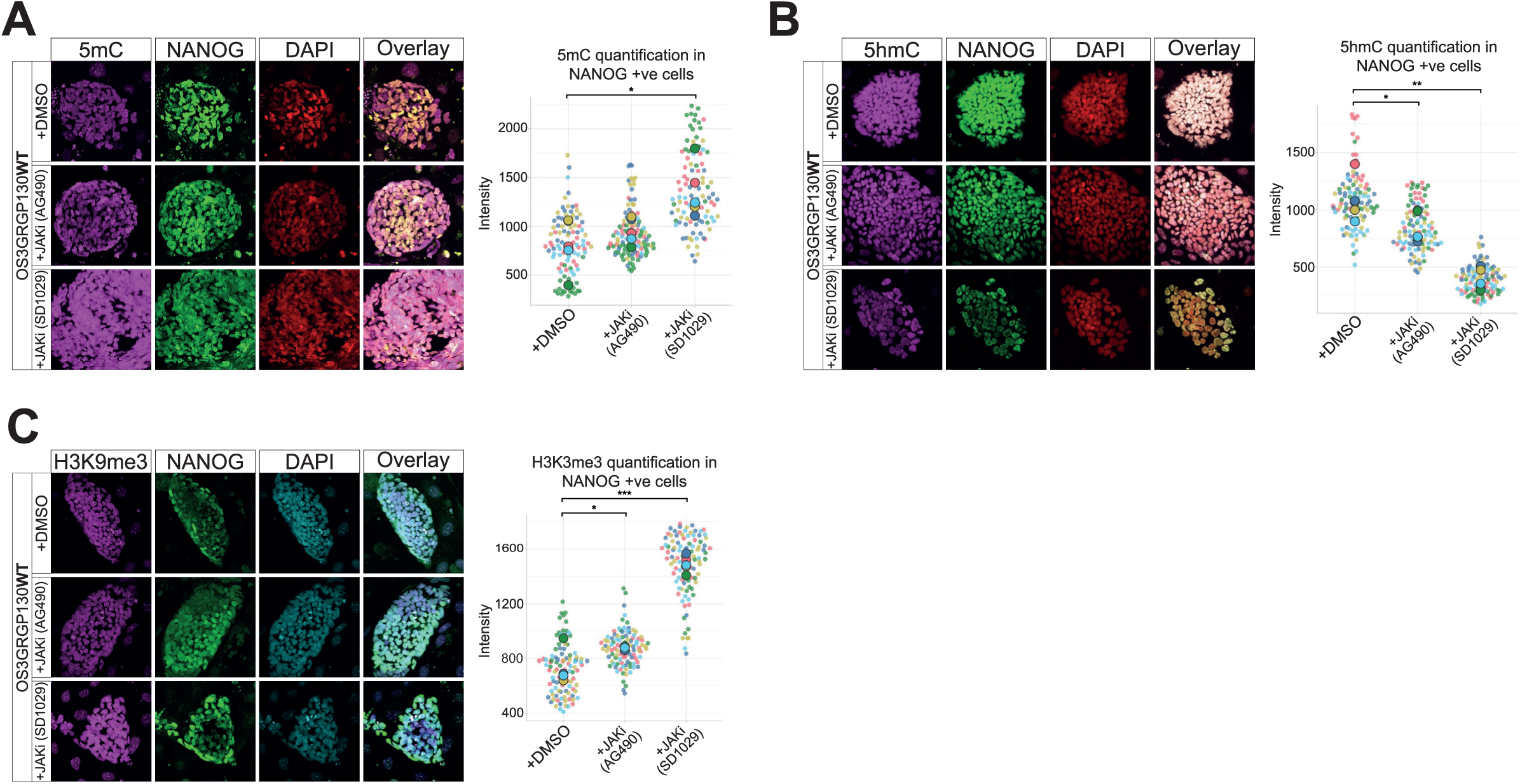
Immunofluorescence Analysis of Epigenetic Marks Following JAK Inhibition. Immunofluorescence labelling of OS3-GRgp130WT cells cultured with 4’OHT+GCSF and JAK inhibitors (AG490 and SD1029), with antibodies to NANOG, (**A**) 5mC, (**B**) 5hmC and (**C**) H3K9me3. The corresponding histograms show the fluorescence quantification in NANOG positive cells. Statistical test: Welch’s *t*-test (* = *p* ≤ 0.05, ** = *p* ≤ 0.01, *** = *p* ≤ 0.001).

**Supplementary figure 6:**
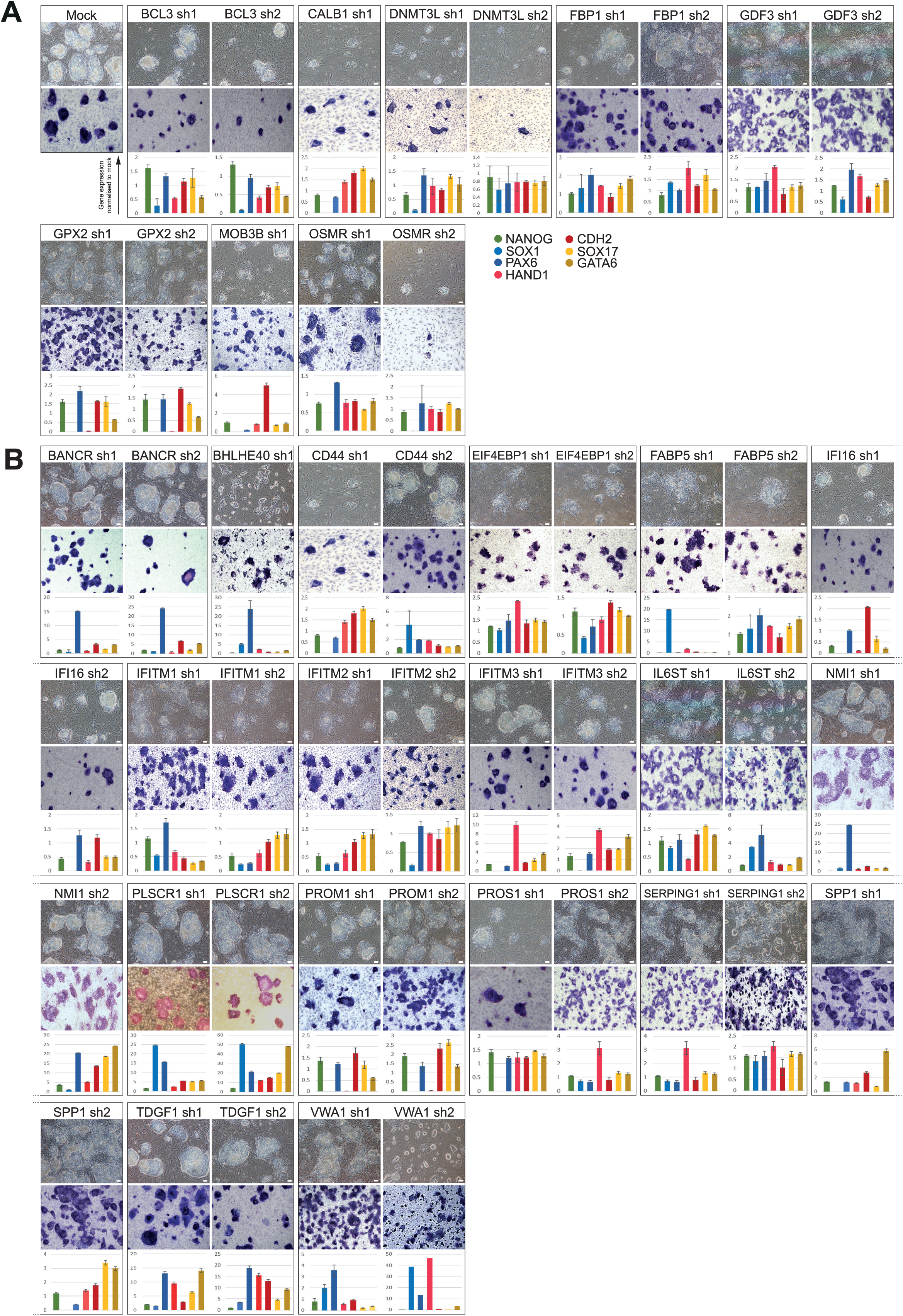
Biological replicate of the analysis presented in. Fig. 5. (**A, B**) Phase-contrast microphotographs, AP activity assay, and qRT-PCR analysis of NANOG, SOX1, PAX6, HAND1, CDH2, SOX17 and GATA6 expression in candidate gene knockdown OS3-GRgp130 cell lines cultured with 4’OHT+GCSF for 7 days.

**Supplementary figure 7:**
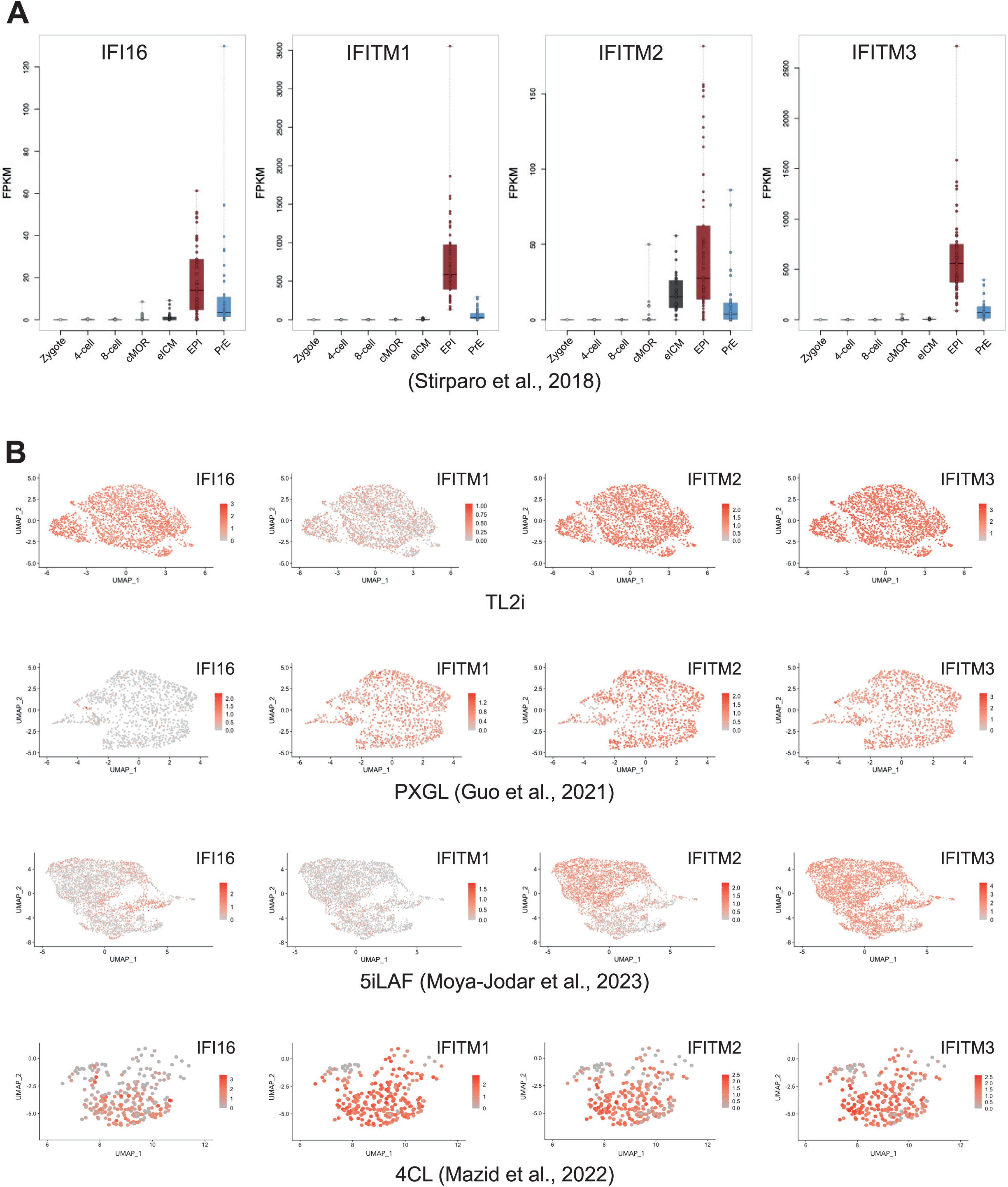
Expression of IFI16 and IFITM Family Genes in Human Pre-implantation Embryos and Naïve PSCs. **(A)** Transcript expression levels (FPKM) of *IFI16*, *IFITM1*, *IFITM2*, and *IFITM3* in human pre-implantation embryos at the following developmental stages: zygote, 4-cell, 8-cell, compacted morula (cMOR), early inner cell mass (eICM), epiblast (EPI), and primitive endoderm (PrE). **(B)** Feature plots showing the expression of *IFI16*, *IFITM1*, *IFITM2*, and *IFITM3* in naïve human pluripotent stem cells (PSCs) reprogrammed under TL2i, PXGL, 5iLAF, and 4CL culture conditions.

**Supplementary figure 8:**
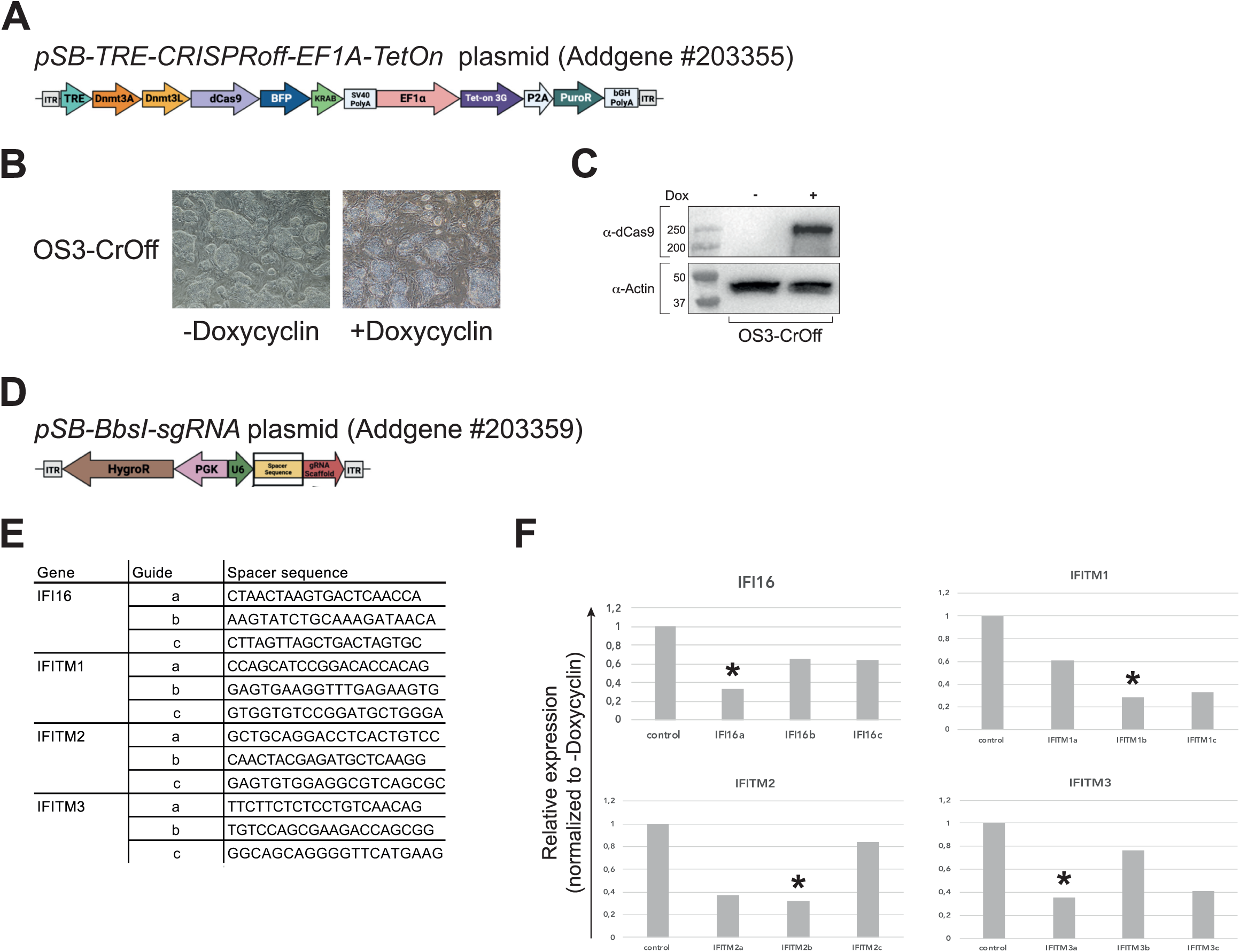
Inducible CRISPRoff System for Targeted Silencing of IFI16 and IFITM Genes. (**A**) Schematic representation of the *pSB-TRE-CRISPRoff-EF1A-TetOn* plasmid (Addgene #203355), enabling doxycycline-inducible expression of dCas9 fused to the DNA methyltransferases DNMT3A and DNMT3L, as well as the KRAB repression domain. (**B**) Phase-contrast micrographs of OS3-CrOff cells cultured in the absence or presence of doxycycline are shown. (**C**) Western blot analysis of dCas9 expression in OS3-CrOff cells under –doxycycline and +doxycycline conditions. (**D**) Schematic representation of the *pSB-BbsI-sgRNA* plasmid (Addgene #203359) driving guide RNA expression. (**E**) Sequences of three sgRNA spacer regions designed to target *IFI16*, *IFITM1*, *IFITM2*, and *IFITM3* are indicated. (F) qRT–PCR analysis of *IFI16*, *IFITM1*, *IFITM2*, and *IFITM3* expression in OS3-CrOff cells following transfection with the corresponding pSB-BbsI-sgRNA plasmids, cultured in the absence or presence of doxycycline. Asterisks (*) indicate the sgRNAs selected for subsequent experiments.

**Supplementary figure 9:**
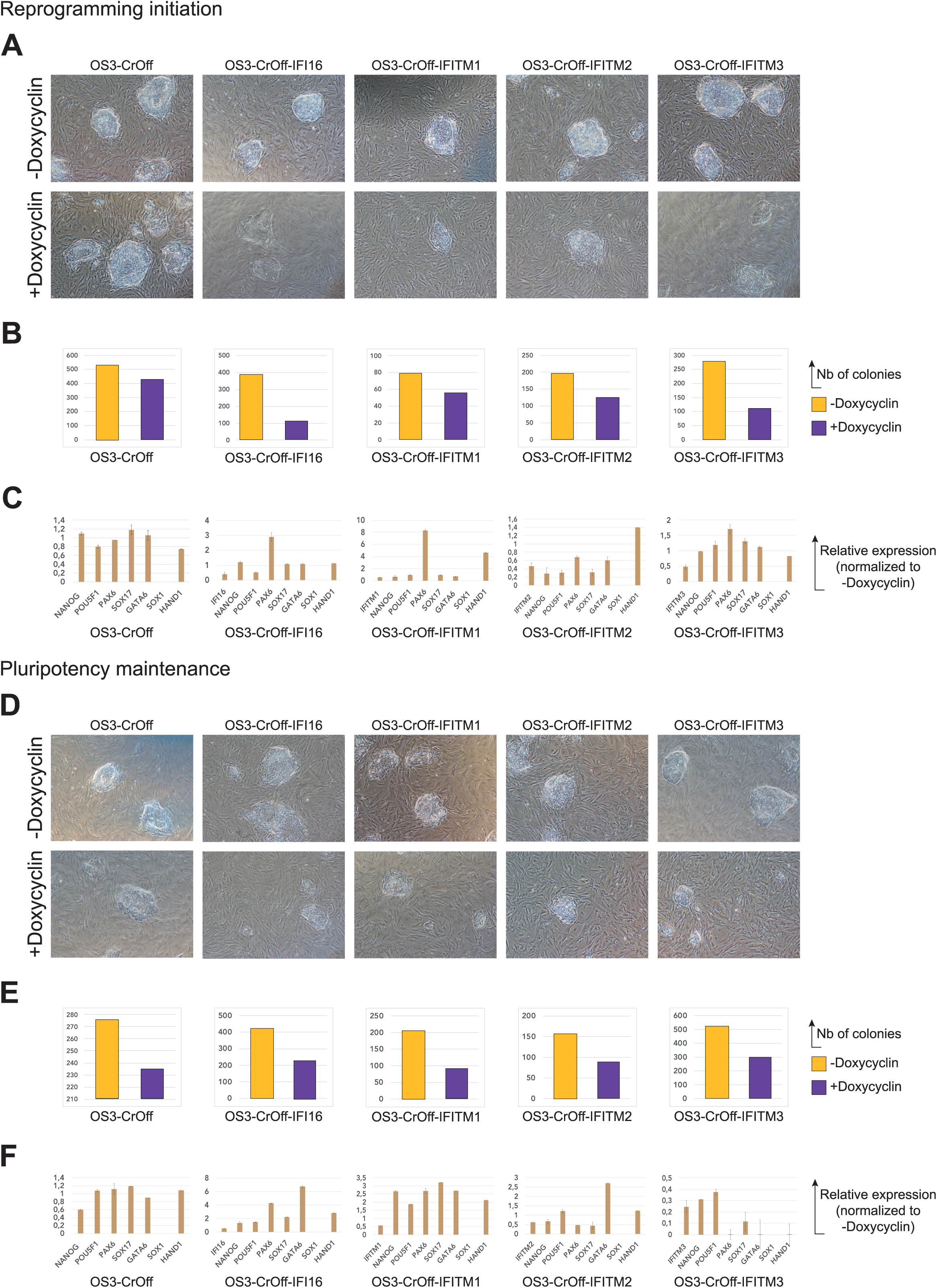
Effects of CRISPRoff-Mediated Gene Silencing on Pluripotency and Differentiation Markers. **(A)** Phase-contrast micrographs of OS3-CrOff, OS3-CrOff-IFI16, OS3-CrOff-IFITM1, OS3-CrOff-IFITM2, and OS3-CrOff-IFITM3 cells plated at clonal density in FGF2-containing medium. FGF2 was subsequently replaced with LIF and tamoxifen, and cells were cultured for seven days in the presence or absence of doxycycline. **(B)** Histogram showing the number of colonies quantified from the alkaline phosphatase (AP) assay presented in Fig. 6A. **(C)** qRT–PCR analysis of pluripotency markers (*NANOG*, *POU5F1*) and lineage-associated genes (*PAX6*, *SOX17*, *GATA6*, *SOX1*, *HAND1*) in OS3-CrOff, OS3-CrOff-IFI16, OS3-CrOff-IFITM1, OS3-CrOff-IFITM2, and OS3-CrOff-IFITM3 cells cultured as described in (A). **(D)** Phase-contrast micrographs of OS3-CrOff, OS3-CrOff-IFI16, OS3-CrOff-IFITM1, OS3-CrOff-IFITM2, and OS3-CrOff-IFITM3 cells plated at clonal density directly in tamoxifen-and LIF-containing medium and cultured for seven days in the presence or absence of doxycycline. **(E)** Histogram showing the number of colonies quantified from the AP assay presented in Fig. 6D. **(F)** qRT–PCR analysis of pluripotency markers (*NANOG*, *POU5F1*) and lineage-associated genes (*PAX6*, *SOX17*, *GATA6*, *SOX1*, *HAND1*) in cells cultured as described in (D).

**Supplementary Table 1.**
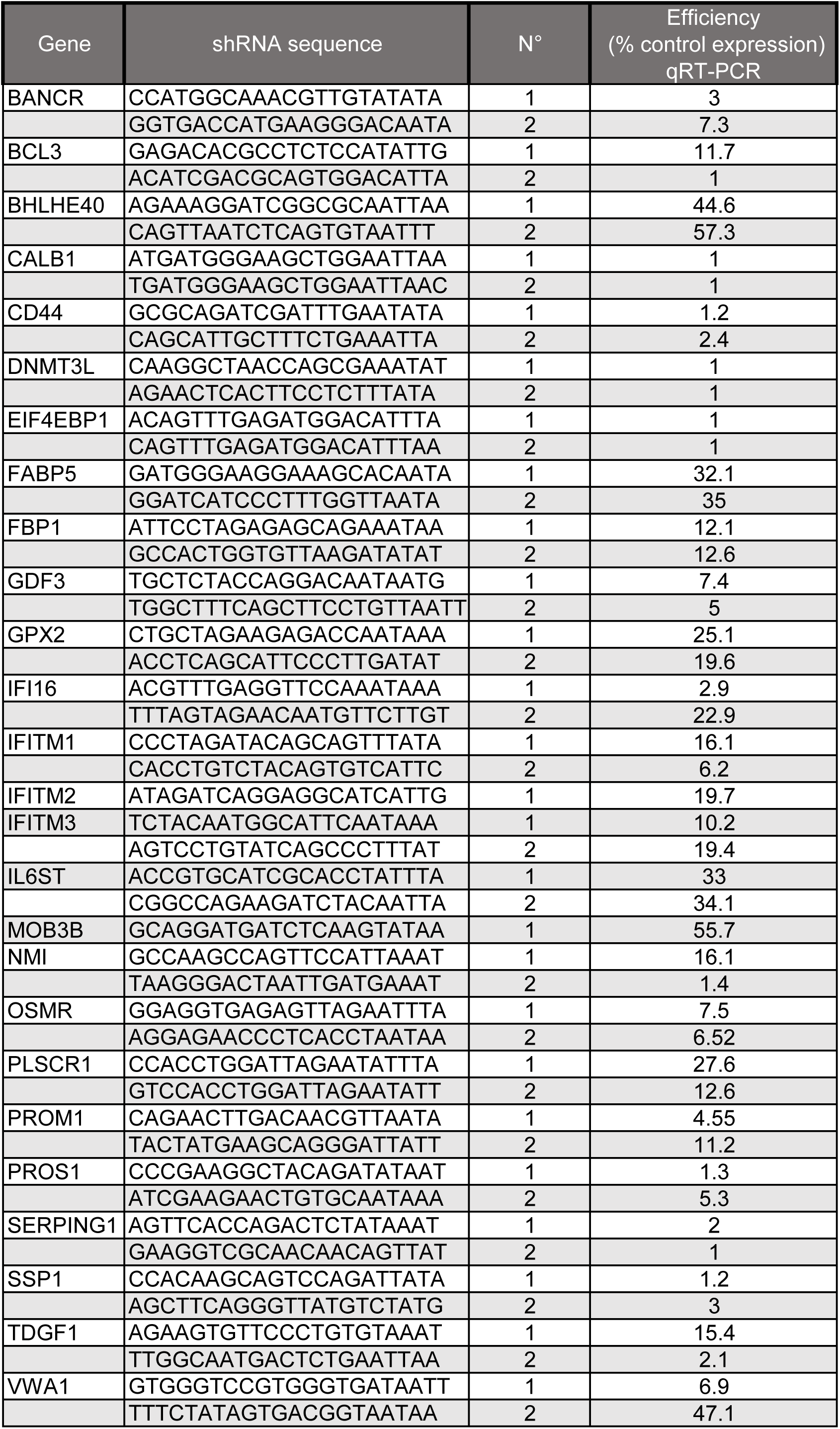

## Literature cited

1. Aksoy I, Giudice V, Delahaye E, Wianny F, Aubry M, Mure M, Chen J, Jauch R, Bogu GK, Nolden T et al (2014) Klf4 and Klf5 differentially inhibit mesoderm and endoderm differentiation in embryonic stem cells. Nat Commun 5: 3719–3733

2. Anneren C, Cowan CA, Melton DA (2004) The Src family of tyrosine kinases is important for embryonic stem cell self-renewal. J Biol Chem 279: 31590–31598

3. Bahn MS, Ko YG (2023) PROM1-mediated cell signal transduction in cancer stem cells and hepatocytes. BMB reports 56: 65–70

4. Bahn MS, Yu DM, Lee M, Jo SJ, Lee JW, Kim HC, Lee H, Kim HL, Kim A, Hong JH et al (2022) Central role of Prominin-1 in lipid rafts during liver regeneration. Nat Commun 13: 6219

5. Boeuf H, Hauss C, Graeve FD, Baran N, Kedinger C (1997) Leukemia inhibitory factor-dependent transcriptional activation in embryonic stem cells. J Cell Biol 138: 1207–1217

6. Bourillot PY, Aksoy I, Schreiber V, Wianny F, Schulz H, Hummel O, Hubner N, Savatier P (2009) Novel STAT3 target genes exert distinct roles in the inhibition of mesoderm and endoderm differentiation in cooperation with Nanog. Stem Cells 27: 1760–1771

7. Bourillot PY, Santamaria C, David L, Savatier P (2019) GP130 signaling and the control of naive pluripotency in humans, monkeys, and pigs. Exp Cell Res: 111712

8. Bowling S, Di Gregorio A, Sancho M, Pozzi S, Aarts M, Signore M, M DS, Martinez-Barbera JP, Gil J, Rodriguez TA (2018) P53 and mTOR signalling determine fitness selection through cell competition during early mouse embryonic development. Nat Commun 9: 1763

9. Burdon T, Stracey C, Chambers I, Nichols J, Smith A (1999) Suppression of SHP-2 and ERK signalling promotes self-renewal of mouse embryonic stem cells. Dev Biol 210: 30–43

10. Cartwright P, McLean C, Sheppard A, Rivett D, Jones K, Dalton S (2005) LIF/STAT3 controls ES cell self-renewal and pluripotency by a Myc-dependent mechanism. Development 132: 885–896

11. Chan YS, Goke J, Ng JH, Lu X, Gonzales KA, Tan CP, Tng WQ, Hong ZZ, Lim YS, Ng HH (2013) Induction of a human pluripotent state with distinct regulatory circuitry that resembles preimplantation epiblast. Cell Stem Cell 13: 663–675

12. Chen H, Aksoy I, Gonnot F, Osteil P, Aubry M, Hamela C, Rognard C, Hochard A, Voisin S, Fontaine E et al (2015) Reinforcement of STAT3 activity reprogrammes human embryonic stem cells to naive-like pluripotency. Nat Commun 6: 7095

13. Daheron L, Opitz SL, Zaehres H, Lensch WM, Andrews PW, Itskovitz-Eldor J, Daley GQ (2004) LIF/STAT3 signaling fails to maintain self-renewal of human embryonic stem cells. Stem Cells 22: 770–778

14. Dal Col J, Lamberti MJ, Nigro A, Casolaro V, Fratta E, Steffan A, Montico B (2022) Phospholipid scramblase 1: a protein with multiple functions via multiple molecular interactors. Cell Commun Signal 20: 78

15. Duggal G, Warrier S, Ghimire S, Broekaert D, Van der Jeught M, Lierman S, Deroo T, Peelman L, Van Soom A, Cornelissen R et al (2015) Alternative Routes to Induce Naive Pluripotency in Human Embryonic Stem Cells. Stem Cells 33: 2686–2698

16. Dutchak K, Garnett S, Nicoll M, de Bruyns A, Dankort D (2022) MOB3A Bypasses BRAF and RAS Oncogene-Induced Senescence by Engaging the Hippo Pathway. Mol Cancer Res 20: 770–781

17. Ernst M, Gearing DP, Dunn AR (1994) Functional and biochemical association of Hck with the LIF/IL-6 receptor signal transducing subunit gp130 in embryonic stem cells. EMBO J 13: 1574–1584

18. Ernst M, Oates A, Dunn AR (1996) Gp130-mediated signal transduction in embryonic stem cells involves activation of Jak and Ras/mitogen-activated protein kinase pathways. J Biol Chem 271: 30136–30143

19. Fiorenzano A, Pascale E, D’Aniello C, Acampora D, Bassalert C, Russo F, Andolfi G, Biffoni M, Francescangeli F, Zeuner A et al (2016) Cripto is essential to capture mouse epiblast stem cell and human embryonic stem cell pluripotency. Nat Commun 7: 12589

20. Friedlova N, Zavadil Kokas F, Hupp TR, Vojtesek B, Nekulova M (2022) IFITM protein regulation and functions: Far beyond the fight against viruses. Front Immunol 13: 1042368

21. Gafni O, Weinberger L, Mansour AA, Manor YS, Chomsky E, Ben-Yosef D, Kalma Y, Viukov S, Maza I, Zviran A et al (2013) Derivation of novel human ground state naive pluripotent stem cells. Nature 504: 282–286

22. Gao X, Nowak-Imialek M, Chen X, Chen D, Herrmann D, Ruan D, Chen ACH, Eckersley-Maslin MA, Ahmad S, Lee YL et al (2019) Establishment of porcine and human expanded potential stem cells. Nat Cell Biol 21: 687–699

23. Geiselmann A, Micouin A, Vandormael-Pournin S, Laville V, Chervova A, Mella S, Navarro P, Cohen-Tannoudji M (2024) PI3K/AKT signaling controls ICM maturation and proper epiblast and primitive endoderm specification in mice. Dev Cell

24. Gil M, Hamann CA, Brunger JM, Gama V (2024) Engineering a CRISPRoff Platform to Modulate Expression of Myeloid Cell Leukemia (MCL-1) in Committed Oligodendrocyte Neural Precursor Cells. Bio Protoc 2024 Jan 5;14(1):e4913

25. Gomez-Herranz M, Taylor J, Sloan RD (2023) IFITM proteins: Understanding their diverse roles in viral infection, cancer, and immunity. J Biol Chem 299: 102741

26. Guo G, Stirparo GG, Strawbridge SE, Spindlow D, Yang J, Clarke J, Dattani A, Yanagida A, Li MA, Myers S et al (2021) Human naive epiblast cells possess unrestricted lineage potential. Cell Stem Cell

27. Guo G, von Meyenn F, Rostovskaya M, Clarke J, Dietmann S, Baker D, Sahakyan A, Myers S, Bertone P, Reik W et al (2017) Epigenetic resetting of human pluripotency. Development 144: 2748–2763

28. Guo G, von Meyenn F, Santos F, Chen Y, Reik W, Bertone P, Smith A, Nichols J (2016) Naive Pluripotent Stem Cells Derived Directly from Isolated Cells of the Human Inner Cell Mass. Stem cell reports 6: 437–446

29. Hanna J, Cheng AW, Saha K, Kim J, Lengner CJ, Soldner F, Cassady JP, Muffat J, Carey BW, Jaenisch R (2010) Human embryonic stem cells with biological and epigenetic characteristics similar to those of mouse ESCs. Proc Natl Acad Sci U S A 107: 9222–9227

30. He Q, Wu Z, Yang W, Jiang D, Hu C, Yang X, Li N, Li F (2020) IFI16 promotes human embryonic stem cell trilineage specification through interaction with p53. NPJ Regen Med 5: 18

31. Hou Y, Wang S, Gao M, Chang J, Sun J, Qin L, Li A, Lv F, Lou J, Zhang Y et al (2021) Interferon-Induced Transmembrane Protein 3 Expression Upregulation Is Involved in Progression of Hepatocellular Carcinoma. Biomed Res Int 2021: 5612138

32. Hussen BM, Azimi T, Abak A, Hidayat HJ, Taheri M, Ghafouri-Fard S (2021) Role of lncRNA BANCR in Human Cancers: An Updated Review. Front Cell Dev Biol 9: 689992

33. Li Y, Rogulski K, Zhou Q, Sims PJ, Prochownik EV (2006) The negative c-Myc target onzin affects proliferation and apoptosis via its obligate interaction with phospholipid scramblase 1. Mol Cell Biol 26: 3401–3413

34. Love MI, Huber W, Anders S (2014) Moderated estimation of fold change and dispersion for RNA-seq data with DESeq2. Genome Biol 15: 550

35. Mak AB, Nixon AM, Kittanakom S, Stewart JM, Chen GI, Curak J, Gingras AC, Mazitschek R, Neel BG, Stagljar I et al (2012) Regulation of CD133 by HDAC6 promotes beta-catenin signaling to suppress cancer cell differentiation. Cell reports 2: 951–963

36. Martello G, Bertone P, Smith A (2013) Identification of the missing pluripotency mediator downstream of leukaemia inhibitory factor. EMBO J 32: 2561–2574

37. Matsuda T, Nakamura T, Nakao K, Arai T, Katsuki M, Heike T, Yokota T (1999) STAT3 activation is sufficient to maintain an undifferentiated state of mouse embryonic stem cells. Embo J 18: 4261–4269

38. Meistermann D, Bruneau A, Loubersac S, Reignier A, Firmin J, Francois-Campion V, Kilens S, Lelievre Y, Lammers J, Feyeux M et al (2021) Integrated pseudotime analysis of human pre-implantation embryo single-cell transcriptomes reveals the dynamics of lineage specification. Cell Stem Cell 28: 1625–1640 e1626

39. Moya-Jodar M, Ullate-Agote A, Barlabe P, Rodriguez-Madoz JR, Abizanda G, Barreda C, Carvajal-Vergara X, Vilas-Zornoza A, Romero JP, Garate L et al (2023) Revealing cell populations catching the early stages of human embryo development in naive pluripotent stem cell cultures. Stem cell reports 18: 64–80

40. Negre D, Mangeot PE, Duisit G, Blanchard S, Vidalain PO, Leissner P, Winter AJ, Rabourdin-Combe C, Mehtali M, Moullier P et al (2000) Characterization of novel safe lentiviral vectors derived from simian immunodeficiency virus (SIVmac251) that efficiently transduce mature human dendritic cells. Gene Ther 7: 1613–1623

41. Niwa H, Burdon T, Chambers I, Smith A (1998) Self-renewal of pluripotent embryonic stem cells is mediated via activation of STAT3. Genes Dev 12: 2048–2060

42. Niwa H, Ogawa K, Shimosato D, Adachi K (2009) A parallel circuit of LIF signalling pathways maintains pluripotency of mouse ES cells. Nature 460: 118–122

43. Paling NR, Wheadon H, Bone HK, Welham MJ (2004) Regulation of embryonic stem cell self-renewal by phosphoinositide 3-kinase-dependent signalling. J Biol Chem 279: 48063–48070

44. Postma M, Goedhart J (2019) PlotsOfData-A web app for visualizing data together with their summaries. PLoS Biol 17: e3000202

45. Prieto AL, Lai C (2024) The TAM Subfamily of Receptor Tyrosine Kinases: The Early Years. Int J Mol Sci 25

46. Qin H, Hejna M, Liu Y, Percharde M, Wossidlo M, Blouin L, Durruthy-Durruthy J, Wong P, Qi Z, Yu J et al (2016) YAP Induces Human Naive Pluripotency. Cell reports 14: 2301–2312

47. Qiu D, Ye S, Ruiz B, Zhou X, Liu D, Zhang Q, Ying QL (2015) Klf2 and Tfcp2l1, Two Wnt/beta-Catenin Targets, Act Synergistically to Induce and Maintain Naive Pluripotency. Stem cell reports 5: 314–322

48. Ricci-Vitiani L, Lombardi DG, Pilozzi E, Biffoni M, Todaro M, Peschle C, De Maria R (2007) Identification and expansion of human colon-cancer-initiating cells. Nature 445: 111–115

49. Sonnentag SJ, Ibrahim NSM, Orian-Rousseau V (2024) CD44: a stemness driver, regulator, and marker-all in one? Stem Cells 42: 1031–1039

50. Storm MP, Bone HK, Beck CG, Bourillot PY, Schreiber V, Damiano T, Nelson A, Savatier P, Welham MJ (2007) Regulation of Nanog expression by phosphoinositide 3-kinase-dependent signalling in murine embryonic stem cells. J Biol Chem 282: 6265–6273

51. Sumi T, Fujimoto Y, Nakatsuji N, Suemori H (2004) STAT3 is dispensable for maintenance of self-renewal in nonhuman primate embryonic stem cells. Stem Cells 22: 861–872

52. Tai CI, Ying QL (2013) Gbx2, a LIF/Stat3 target, promotes reprogramming to and retention of the pluripotent ground state. J Cell Sci 126: 1093–1098

53. Takashima Y, Guo G, Loos R, Nichols J, Ficz G, Krueger F, Oxley D, Santos F, Clarke J, Mansfield W et al (2014) Resetting Transcription Factor Control Circuitry toward Ground-State Pluripotency in Human. Cell 158: 1254–1269

54. Tanaka SS, Yamaguchi YL, Tsoi B, Lickert H, Tam PP (2005) IFITM/Mil/fragilis family proteins IFITM1 and IFITM3 play distinct roles in mouse primordial germ cell homing and repulsion. Dev Cell 9: 745–756

55. Taniguchi K, Wu LW, Grivennikov SI, de Jong PR, Lian I, Yu FX, Wang K, Ho SB, Boland BS, Chang JT et al (2015) A gp130-Src-YAP module links inflammation to epithelial regeneration. Nature 519: 57–62

56. Theunissen TW, Powell BE, Wang H, Mitalipova M, Faddah DA, Reddy J, Fan ZP, Maetzel D, Ganz K, Shi L et al (2014) Systematic Identification of Culture Conditions for Induction and Maintenance of Naive Human Pluripotency. Cell Stem Cell 15: 471–487

57. Wang H, Tang F, Bian E, Zhang Y, Ji X, Yang Z, Zhao B (2020) IFITM3/STAT3 axis promotes glioma cells invasion and is modulated by TGF-beta. Mol Biol Rep 47: 433–441

58. Wang W, Lin C, Lu D, Ning Z, Cox T, Melvin D, Wang X, Bradley A, Liu P (2008) Chromosomal transposition of PiggyBac in mouse embryonic stem cells. Proc Natl Acad Sci U S A 105: 9290–9295

59. Wang ZY, Wang PG, An J (2021) The Multifaceted Roles of TAM Receptors during Viral Infection. Virol Sin 36: 1–12

60. Ward AC, Monkhouse JL, Csar XF, Touw IP, Bello PA (1998) The Src-like tyrosine kinase Hck is activated by granulocyte colony-stimulating factor (G-CSF) and docks to the activated G-CSF receptor. Biochem Biophys Res Commun 251: 117–123

61. Ware CB, Nelson AM, Mecham B, Hesson J, Zhou W, Jonlin EC, Jimenez-Caliani AJ, Deng X, Cavanaugh C, Cook S et al (2014) Derivation of naive human embryonic stem cells. Proc Natl Acad Sci U S A 111: 4484–4489

62. Wu B, Li Y, Li B, Zhang B, Wang Y, Li L, Gao J, Fu Y, Li S, Chen C et al (2021) DNMTs Play an Important Role in Maintaining the Pluripotency of Leukemia Inhibitory Factor-Dependent Embryonic Stem Cells. Stem cell reports 16: 582–596

63. Wulansari N, Sulistio YA, Darsono WHW, Kim CH, Lee SH (2021) LIF Maintains mESC Pluripotency by Modulating TET1 and JMJD2 Activity in a JAK2-Dependent Manner. Stem Cells 39

64. Yang G, Xu Y, Chen X, Hu G (2007) IFITM1 plays an essential role in the antiproliferative action of interferon-gamma. Oncogene 26: 594–603

65. Yang Y, Liu B, Xu J, Wang J, Wu J, Shi C, Xu Y, Dong J, Wang C, Lai W et al (2017) Derivation of Pluripotent Stem Cells with In Vivo Embryonic and Extraembryonic Potency. Cell 169: 243–257 e225

66. Ye X, Tian C, Liu L, Feng G, Jin K, Wang H, Chen J, Liu L (2021) Oncostatin M Maintains Naive Pluripotency of mESCs by Tetraploid Embryo Complementation (TEC) Assay. Front Cell Dev Biol 9: 675411

67. Zhang G, Xie Y, Zhou Y, Xiang C, Chen L, Zhang C, Hou X, Chen J, Zong H, Liu G (2017) p53 pathway is involved in cell competition during mouse embryogenesis. Proc Natl Acad Sci U S A 114: 498–503

68. Zhao B, Wang H, Zong G, Li P (2013) The role of IFITM3 in the growth and migration of human glioma cells. BMC Neurol 13: 210

69. Zhao H, Gonzalezgugel E, Cheng L, Richbourgh B, Nie L, Liu C (2015) The roles of interferon-inducible p200 family members IFI16 and p204 in innate immune responses, cell differentiation and proliferation. Genes Dis 2: 46–56

